# The cellular associates of late life changes in white matter microstructure

**DOI:** 10.1101/2025.09.02.673734

**Authors:** Rowena Chin, Xi-Han Zhang, Kevin M. Anderson, Anastasia Yendiki, Avram J. Holmes

## Abstract

The microstructural architecture of white matter supporting information flow across local circuits and large-scale networks changes throughout the lifespan. However, the genetic and cellular factors underlying age-related variations in white matter microstructure have yet to be established. Here, we examined the genetic associates of individual differences in diffusion-based measures of white matter in a population-based cohort (N=29,862) from the UK Biobank. Estimates of heritability from Genome-Wide Association Study (GWAS) data revealed that genetic factors are linked to population variability in 96.1% of 432 tract microstructural measures. The presence of shared genetic influences was observed to be greater within, relative to between, broad tract classes (commissural, association, projection, and complex cerebellar). Age associations with microstructural changes were estimated across diffusivity measures, with association class tracts showing the greatest vulnerability to age-related decline in older adults. Analyses of imputed cellular associates of age-related changes in white matter revealed a preferential relationship with cell gene markers of oligodendrocytes and other glial cell types, with sparse relationships observed for inhibitory and excitatory cells. These data indicate that white matter tract microstructure is shaped by genetic factors and suggest a role for glial cell-related transcripts in late-life changes in the structural wiring properties of the human brain.

**Significance Statement:** The structural wiring of the human brain supports interregional communication and is essential for cognition, yet the genetic and cellular drivers underlying age-related changes are not well understood. Here, we characterize the extent to which genetic factors follow the major organizational patterns of white matter connectivity and associated age-related changes in late life. Age-related changes were most pronounced in late-developing association tracts, key pathways for higher cognitive functions. By integrating genetic and transcriptomic data, we show that genetic variation linked to glial cell markers accounts for a substantial proportion of age-related white matter changes. These findings provide a mechanistic framework for understanding genetic and cellular determinants of structural brain aging.

## Introduction

Information processing in the human brain is supported by a complex mosaic of spatially distributed functional parcels connected by anatomical tracts (1, 2). These white matter connections facilitate signal propagation across distal brain regions and, when impaired, are linked to heritable psychiatric and neurological disorders, including late-life cognitive decline (3–5). Although large-scale fiber bundles consist of axonal projections of pyramidal cells, they are further supported by the coordinated functions of non-neuronal cells. For instance, glial and their associated trophic factors support axonal myelination and the health of projection cell bodies (6). Critically, white matter tracts are not static across the lifespan and undergo a series of stereotyped structural and functional changes in late life (7–9). However, while aging related changes in white matter emerge from the convergence of both heritable and environmental influences, the cellular associates of late-life alterations in structural connectivity remain to be established.

Individual variation in white matter is linked to a complex genetic architecture (10, 11). Recent twin and population-based studies indicate that *in vivo* diffusion-based (dMRI) estimates of white matter microstructure demonstrate moderate to high heritability, with estimates varying across age groups (10–16). Genome-wide association studies (GWAS) have nominated genetic loci associated with dMRI-derived white matter microstructural phenotypes (11, 17, 18), however the extent to which specific cell-types and associated cellular processes may underpin population-level variability in brain microstructure and connectivity has yet to be systematically investigated. On a cellular level, white matter largely consists of axons (myelinated and unmyelinated), glial cells such as myelin-producing oligodendrocytes, astrocytes, microglia, oligodendrocyte progenitor cells (OPCs) (19), and interstitial neurons dispersed amongst fibers (20). Connectomic studies in the human and non-human mammalian brain have also revealed that cortical regions that are structurally connected by axon projections are more likely to share similar features in cytoarchitecture, such as neuronal density, laminar differentiation, and profiles of gene expression (21–25). Moreover, cytoarchitectural similarities among connected brain regions likely reflect greater levels of coordinated cortical maturation (26, 27). Establishing the cellular associates of *in vivo* population-level variability in white matter tract microstructure could provide insight into the biological underpinnings of dMRI-based white matter indices and structural brain alterations across the lifespan.

Cerebral white matter aging is characterized by a multitude of neural hallmarks such as loss of myelination (28), axonal degeneration (29), increased inflammation (30), and accelerated volumetric loss, even after accounting for the presence of gray matter atrophy (31). On a macroscale level, aspects of white matter microstructure captured by dMRI have revealed common age associated alterations, such as an overall decrease in microstructural anisotropy and increased mean diffusivity observed across white matter tracts (32). There is unequivocal support for white matter changes in aging, with prior work demonstrating both antero-posterior (33, 34) and superior-inferior (35, 36) gradients of age-related decline. Embedded within these gradient transitions, preferential aging effects have been observed within broad tract classes defined through their shared patterns of connectivity, such as association, prefrontal callosal, and prefrontal projection tracts (37–39). This topographic pattern of white matter decline is theorized to reflect an inversion of the sequence of myelination that characterizes early development (40, 41), lending support to the “last-in-first-out” theory of retrogenesis across the lifespan (24, 42–44). Here, tracts that reach maturation last tend to be most vulnerable to aging processes (42). Despite population-level variability in the presence of these aging trajectories, the cellular and genetic bases underlying this scheduled pattern of age-related decline have not been systematically investigated.

In the present study, we examine the association between dMRI-based estimates of tract microstructure, the spatial distribution of cortical cell types, inferred from patterns of gene transcription in bulk tissue data, and age-related changes in white matter. First, in a large population-based sample (N=29,862) (45), we confirm that measures of white matter tract microstructure are under genetic control through estimation of SNP-based heritability from GWAS analyses. In doing so, we establish the presence of increased genetic similarity for white matter tract phenotypes within, relative to between, broad tract classes. Cell-type imputation analyses revealed that genetic variation among cell gene markers for oligodendrocytes and other glial cell types explained an enriched proportion of heritable variance in age-related microstructural changes. These data help establish the cellular and genetic underpinnings of the wiring properties of the human brain, suggesting the presence of associations linking specific cell-types with age-related changes in the white matter.

## Results

### Heritability of white matter tract measures

Population-level variability in the integrity of white matter tracts is under strong genetic control (10, 11, 18, 46). While the associated genetic factors are complex and distributed across the genome, they may aggregate within gene sets that are preferentially expressed within specific tissues and cell types (47), providing information on the cellular associates of white matter tract integrity across the lifespan. Here, we investigated if single-nucleotide polymorphisms (SNPs) that explain heritable variance in diffusion-based estimates of white matter microstructural changes in aging are preferentially enriched within genes linked to particular cell types. To that end, we first sought to establish the genome-wide heritability underlying measures of diffusion MRI microstructure.

Using linkage-disequilibrium score regression (LDSC) and estimates of SNP-based heritability (*h^2^*) from GWAS analyses, we estimated the extent to which population-level variability in white matter can be explained by common genetic variation. Individual-specific diffusion metrics for each participant (n=29,942) were obtained from the UK Biobank (45, 48). Analyses considered 9 white matter microstructural measures for 48 tracts, including diffusion tensor imaging (DTI) measures such as fractional anisotropy (FA), mean diffusivity (MD), L1–L3, mode of anisotropy (MO), as well as neurite orientation dispersion and density imaging (NODDI) measures such as intracellular fluid volume (ICVF), isotropic volume fraction (ISOVF), and orientation dispersion index (OD). While FA and MD are commonly reported measures that capture the coherence and magnitude of diffusion, they cannot reliably differentiate between neural tissue compartments. Measures defined by NODDI may offer a complementary and more mechanistic account of white matter aging. Expanded explanations of these microstructural measures are provided in the Supplementary Information.

Overall, genetic factors accounted for a modest portion of the variance of the microstructural parameters in all white matter tracts. Of the 432 tract measures, 415 (96.1%) were significantly heritable (Bonferroni corrected p<0.05, testing for 432 measures), ranging from 4.4 to 53.4% of the variance (mean *h^2^*=28.5%; Figure 1A, Supplementary Table 1). Grouped across diffusivity indices, mean *h^2^* for 48 individual white matter tracts was 34.5% for FA, 26.2% for MD, 27.3% for L1, 29.3% for L2, 29.2% for L3, 24.7% for MO, 40.7% for ICVF, 18.8% for ISOVF, and 25.7% for OD (Table S1). Homologous tracts of the left and right hemispheres showed comparatively similar heritability across the 9 diffusivity measures (range of Pearson’s *r_s_*=0.81-0.99, p_s_= 8.4e-06-3.4e-16; Supplementary Figure 1).

**Figure 1.**
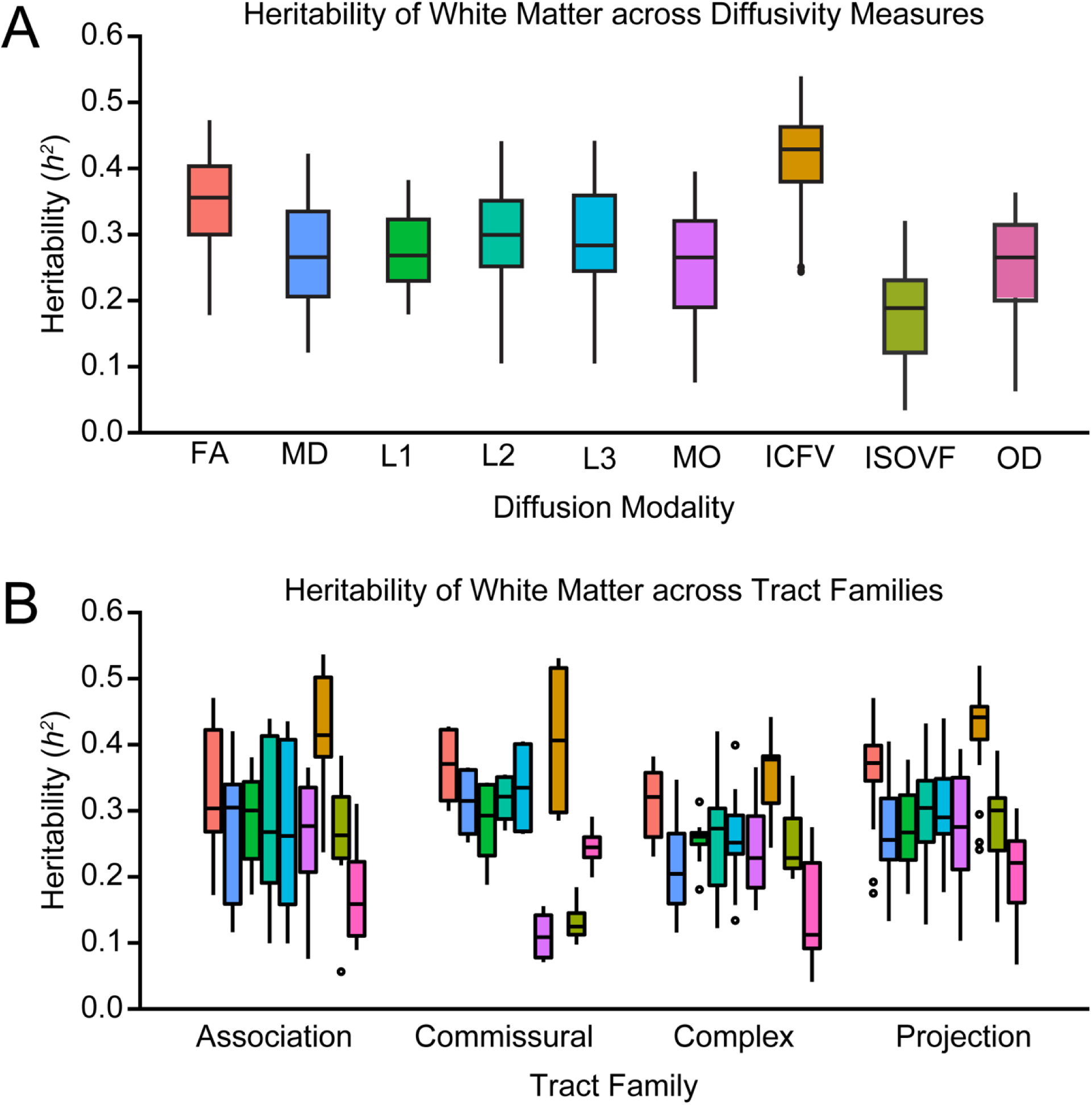
SNP-based heritability estimates across diffusivity metrics in 29,862 participants. (A) SNP-based heritability analyses across 48 white matter tracts labeled according to the JHU ICBM-DTI-81 White Matter Atlas (122) across 9 diffusivity modalities (432 diffusivity-based tract measures). Boxplots are colored according to diffusivity modality and show median as the thick black line; end points of the box mark 25th and 75^th^ percentiles. Across tract measures, 96.1% showed significant SNP-based heritability (p<0.05) after Bonferroni correction, ranging from 4.4% to 53.4%. (B) SNP-based heritability measures are categorized into 4 main white matter tract classes, namely Association, Commissural, Complex, and Projection.

White matter tracts are often organized into groups based on the location of the regions that they connect (49–51). Accordingly, we categorized tracts into four primary groups: complex, associative, commissural, and projection tract fibers (52) (See Supplementary Table 2 for a complete classification). A similar categorization framework was also utilized in prior work probing the genetic bases of white matter tract communities and genetic pleiotropy with cognitive and mental health traits (11). When grouped into 4 tract classes, we did not observe evidence for differences in SNP heritability (F(3,428)=0.564, p=0.639; Figure 1B), where mean *h^2^*was 28.5% in association, 28.6% in commissural, 27.2% in complex, and 29.0% in projection tracts. These data support broad consistency in the proportion of the phenotypic variation that can be explained by genetic factors across tract classes.

### Genetic similarity between white matter tract phenotypes within tract classes

The functional and computational capabilities of brain areas are determined not only by their local properties, but also by their connectivity profiles (53, 54). Converging evidence from *ex-vivo* work in human and non-human animal models have allowed for broad classifications of major white matter bundles according to aspects of their anatomical connectivity (49, 50, 55). For instance, locational features such as the regions that a given set of tract connects to and/or traverses. Intriguingly, tracts that are broadly grouped based on their connectivity patterns possess common developmental and lifespan trajectories (9, 41), susceptibility to aging processes (56), and disease related disruption (for example, in Alzheimer’s disease; (57)).

However, the extent to which common or distinct genetic factors may associate with microstructural features of these broad tract classes remains to be determined. Accordingly, we next examined the extent to which patterns of genetic similarity may be more similar between white matter tract phenotypes of the same tract class compared to tracts from another class.

To examine the shared or distinct genetic factors associated with tract integrity, we estimated genetic correlation (r_G_) for pairs of white matter tract phenotypes across the 9 microstructural measures using the Genome-wide Complex Trait Analysis (GCTA) bivariate genomic-relatedness-based restricted maximum likelihood (GREML) utility. This approach captures SNP-based genetic variance explained by phenotypic correlations between traits (see Methods). These analyses yielded genetic correlational matrices for each microstructural measure that were constructed based on gene correlation (r_G_) estimates between whole brain white matter tracts, arranged according to the four respective tract classes (Figure 2 and Supplementary Data).

**Fig 2.**
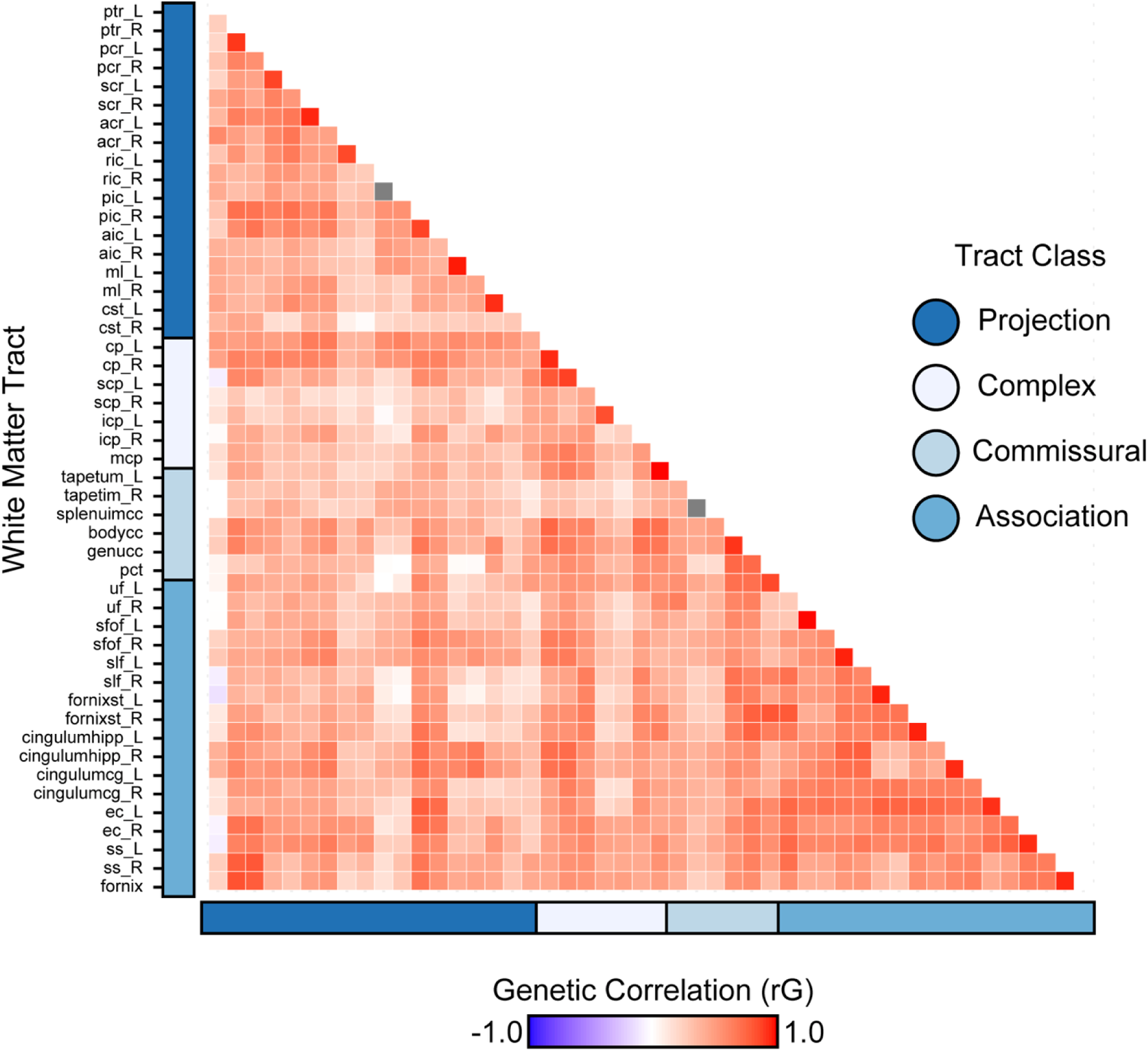
Gene correlation map of whole brain white matter tract phenotypes as estimated by fractional anisotropy (FA). Genetic correlation scores (r_G_) were estimated across all single-nucleotide polymorphisms (SNPs) for 48 pairs of white matter tract phenotypes using LD-Score regression, for FA. A definition of tract abbreviations is provided in the Supplementary Table S1. Analyses were repeated across 9 diffusivity modalities (see Supplement). Within the gene correlational matrix, white matter tracts were grouped according to one of four respective tract classes: Association, Commissural, Complex, Projection (additional details available in Supplementary Table S2). Stronger positive correlations (indicated in red) reflect higher genetic similarity between white matter tract phenotypes, with weaker correlations depicted at lower color intensity.

When considering FA, gene correlation estimates (r_G_) were significantly higher across white matter tract phenotypes within the same tract class in comparison to tracts from other distinct classes (Figure 3). In particular, r_G_ estimates were significantly higher within-than between-tract classes for association (t=2.36, p=0.019), projection (t=13.37, p<2.2e-16), commissural (t=2.68, p=0.020), and complex cerebellar (t=2.78, p=0.011), suggesting white matter tracts within their respective classes generally share greater genetic similarity than between other tract classes. Cross-examination of other related microstructural measures such as MD showed similar patterns, with r_G_ estimates that were significantly higher within-than between-tract classes for association (t=5.88, p=1.875e-08, projection (t=3.61, p=3.834e-4), and complex cerebellar (t=4.26, p=3.265e-4), all other tract class comparisons (p’s>0.05; full results available in Supplemental Table 4). In addition, significant within- and between-tract class differences in r_G_ estimates for NODDI measures were also present across association (ICVF: t=6.97, p=5.5e-11; ISOVF: t=4.19, p=4.2e-05; OD: t=5.44, p=1.9e-07), projection (ICVF: t=6.40, p=1.1e-09; ISOVF: t=8.79, p=1.4e-15; OD: t=3.36, p=0.001), commissural (ISOVF: t=2.20, p=0.047; OD: t=2.24, p=0.044), and complex cerebellar tracts (ICVF: t=2.28, p=0.033; ISOVF: t=5.65, p=1.3e-05; OD: t=4.20, p<0.001). Taken together, these data demonstrate increased genetic similarity across white matter tracts within tract classes than tracts between classes.

**Figure 3.**
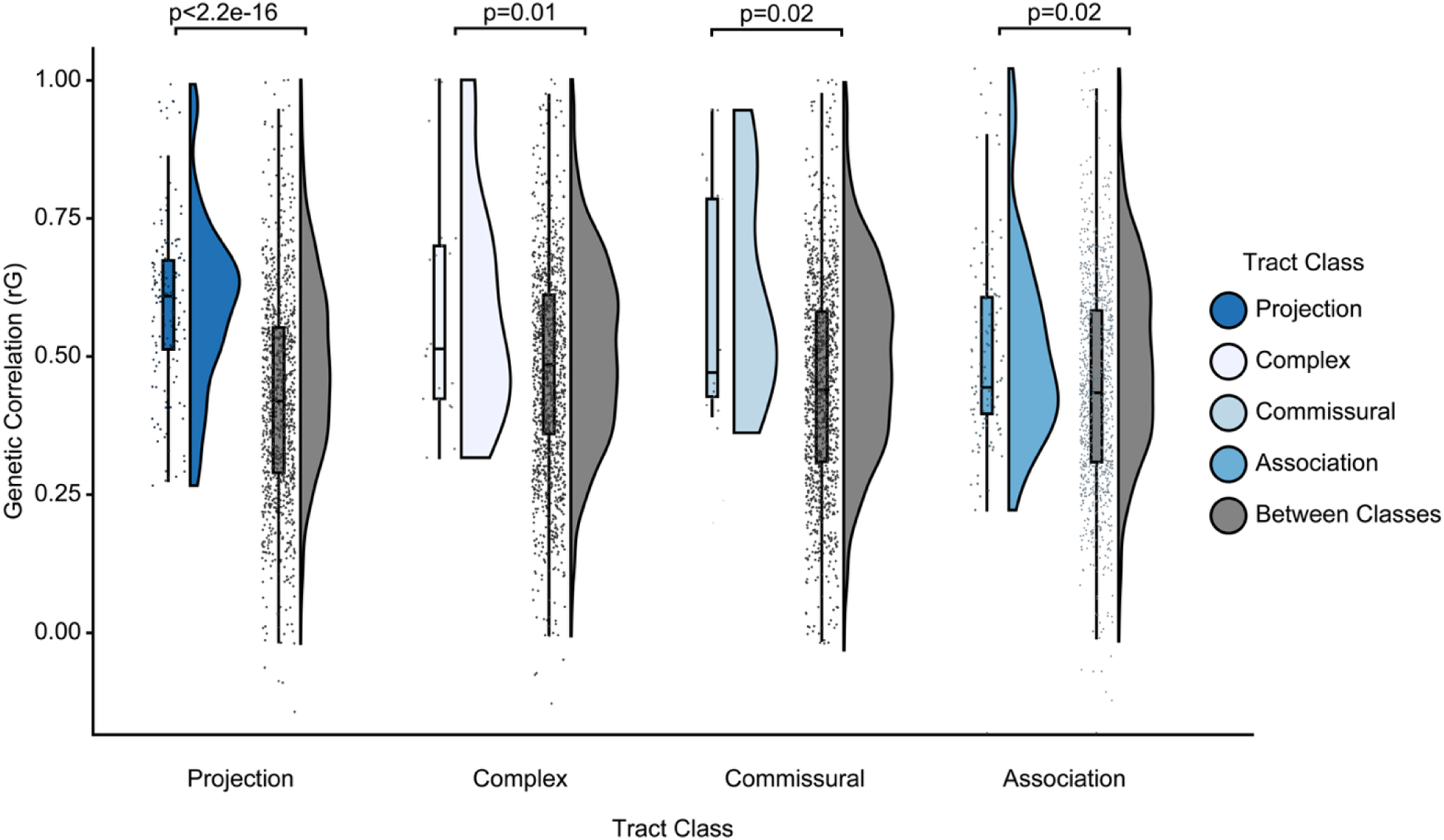
Genetic similarity is greater within white matter tract classes than between tract classes. When considering FA, estimated genetic correlation scores (r_G_) were significantly higher across white matter tract phenotypes within the same tract class compared to tracts from another class, within- and between-association, projection, and complex tracts were quantitatively compared (p<0.05). Violin plots are colored according to white matter tract class (within-tract comparisons are depicted in blue shades and between-tract comparisons are in gray) and show median as the thick black line; end points of the box mark 25th and 75^th^ percentiles. Scatter plots show the relative spread of all pair-wise comparisons between tracts. Analyses of other diffusivity modalities are available in the supplement.

### Age-related white matter changes

At the population-level, late-life aging is associated with a reduction in global brain volumes (58, 59), thinning in cerebral cortical gray matter (58, 60, 61), and increased ventricle sizes (62, 63), along with wide-ranging alterations in white matter including atrophy and lesions (61, 64–66), reduced tract integrity (33, 67), inflammation (68, 69), and a loss of myelination (28, 70). Consistent with prior work on aging-related changes in white matter (11, 37) in the present data, older age was associated with lower coherence of water diffusion (FA; β’s≥-0.436), lower neurite density (ICVF; β’s≥-0.284), lower fiber dispersion (OD; β’s≥-0.167), and conversely, with higher diffusivity (MD; β’s≤0.426) and contribution of isotropic diffusion (ISOVF; β’s≤0.476) across white matter tracts (refer to Supplementary Data S11). This pattern was broadly evident across diffusion measures, indicating a decline in the health of white matter microstructure with older age. In addition, diffusion tensor eigenvalues, i.e., axial diffusivity /L1 (λ1; β’s≤0.308), L2 (λ2; β’s≤0.413) and L3 (λ3; β’s≤0.414) that are components of radial diffusivity, show comparable age associations to MD across tracts. Mode of anisotropy (MO), which is sensitive to crossing fibers, was associated both positively and negatively with age across tracts (-0.322≤b’s≥0.319). The tissue properties reflected in MO estimates remain to be fully determined (71, 72). Although speculative, the present results may reflect alterations in the complexity of white matter architecture and organizational properties as reflected in spatial variability in crossing fibers within across voxels (71).

Association tracts, relative to other tract classes, are more vulnerable to age-related decline (37, 73). In line with this prior work, we observed that the magnitude of associations with age were significantly greater for association class tracts than those of the projection (t=-2.72, p=0.010) and complex cerebellar class tracts in FA (t=-2.21, p=0.038; Figure 4B; see Supplemental for full range of results). Across a range of microstructural indices, significant differences in age-related associations between association class tracts and other tract classes were observed, such as for MD and ICVF, where association tracts also showed greater decline than projection (MD: t=9.91, p<2.2e-16; ICVF: t=-2.63, p=0.014) and complex cerebellar tracts (MD: t=22.29, p<2.2e-16; ICVF: t=-7.45, p=2.7e-06 for association vs complex tracts), and for MD and OD, where association tracts displayed greater decline than commissural tracts (MD: t=-3.13, p=0.002; OD: t=2.61, p=0.042). There were no significant differences between tract classes for measures of ISOVF (p’s>0.05). Aging effects were marked in association tracts such as the fornix and fornix stria terminalis, which exhibited the highest magnitude of age-related changes across 8 out of 9 microstructural measures (refer to Supplementary Data S11). Highlighting the bilateral nature of the observed late-life changes, consistent age-related microstructural profiles were evident within homologous tracts across hemispheres (*r*(187)=0.976, p<2.2e-16). Whole brain white matter spatial maps of age association across tracts for FA are displayed in Figure 4A, as well as presented in Supplementary Figure 2 for the full set of microstructural measures.

**Figure 4.**
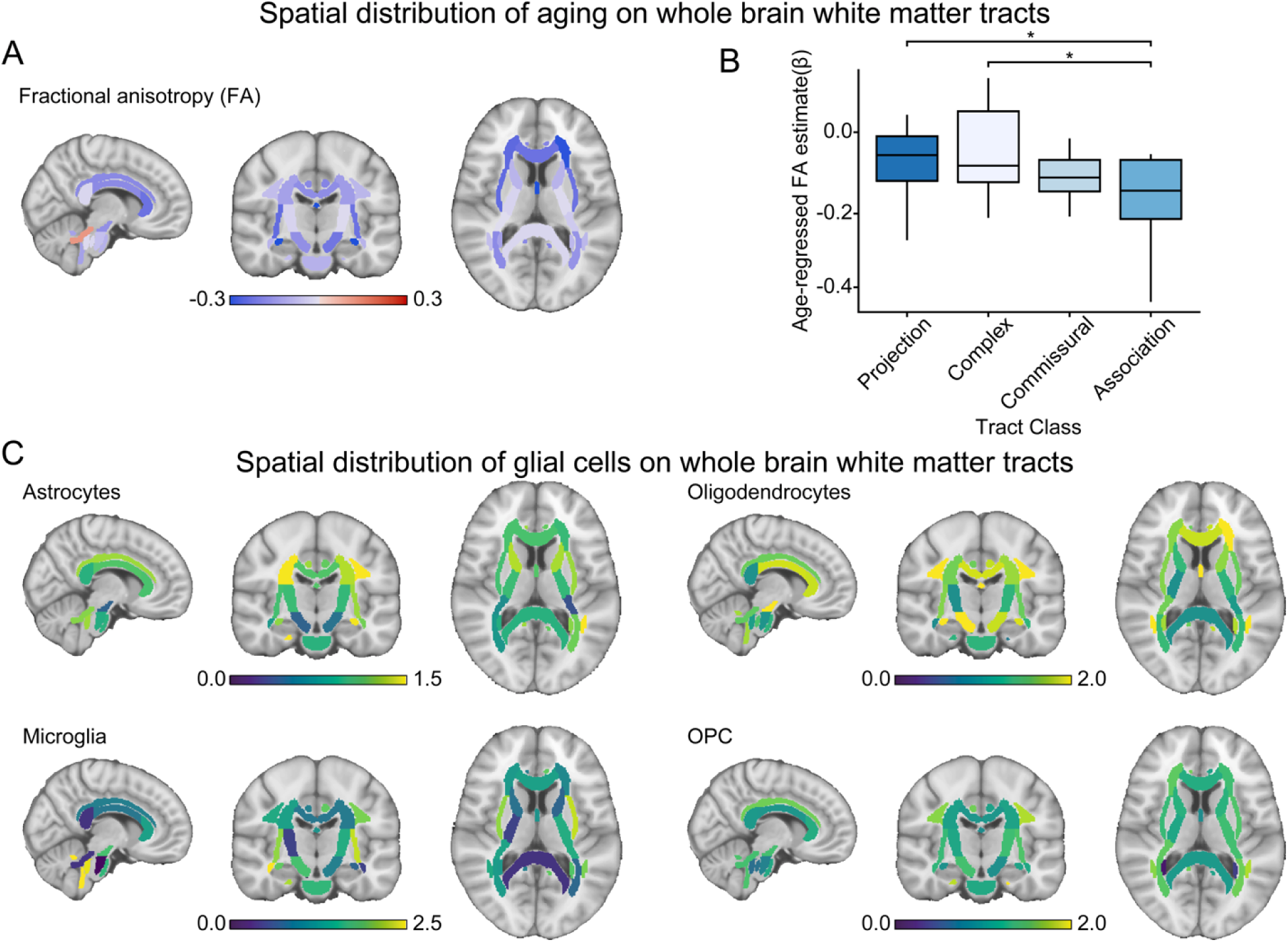
Spatial distribution of age associations and glial cell enrichment across whole brain white matter indexed by fractional anisotropy (FA). (A) Spatial distribution of age associations across whole brain white matter tracts as measured by fractional anisotropy (FA) depict a significant negative age association with FA (blue indicates negative association, red indicates positive association). (B) Age-regressed FA estimates (b) show significant differences between white matter tract classes (p<0.05). Association tracts generally show the greatest negative age-regressed FA, significantly greater than projection (t=-2.72, p=0.010) and complex cerebellar tracts (t=-2.21, p=0.038). All reported p-values are FDR-corrected. (C) Spatial distribution of glial cell enrichment (i.e., astrocytes, microglia, oligodendrocytes, OPC) on whole brain white matter FA tract measures estimated using LDSC partitioned heritability (Dark blue indicates low enrichment, bright yellow indicates high enrichment). Analyses of other diffusivity modalities are available in the supplement.

### Cellular associates of aging in whole brain white matter tracts

Although the polygenic architecture of brain white matter has been broadly explored through large-scale genome wide association studies (GWAS) (10, 11, 46, 74, 75), the extent to which cellular profiles are associated with major white matter connections of the brain has yet to be fully characterized *in vivo*. Furthermore, despite extensively documented observations of aging effects on microstructural properties of white matter (5, 7, 8, 34, 37, 73, 76, 77), an important question still remains if spatial distributions of nominated genes that are linked to cell type profiles track respective age-associated brain white matter microstructural changes.

Next, we tested whether SNPs proximal to coding regions of genes that show preferential expression to respective cell types would explain more genetic variance than would be expected by chance, and if this would be associated with age-related alterations in brain white matter measures. This was done by utilizing differentially expressed enriched gene markers (snDrop-seq) from post-mortem bulk tissue samples aggregated across frontal and visual cortices to derive the molecular signature profiles of 20 cell classes (see Methods) (78). This comprised 7 excitatory neuron subtypes (i.e., Ex1, Ex2, Ex3, Ex4, Ex5, Ex6, and Ex8), 6 interneuron subtypes (i.e., In1, In2, In3, In4, In6, In7), and 8 non-neuronal subtypes (i.e., astrocytes, endothelial, microglia, oligodendrocytes, oligodendrocyte precursor cells, pericytes, and Purkinje cells). Enrichment estimates (*h*^2^/snp) were calculated as the proportion of heritability explained by the SNP partition, divided by the fraction of SNPs in that partition. Spatial maps of whole brain white matter of cell type enrichment are provided in Supplementary Figures 3-11, with detailed results available in Supplementary Data S12-20. Cell type enrichment estimates were then examined relative to age-associated microstructural alterations in whole brain white matter tracts. Statistical association was calculated between aging correlates of 48 whole brain white matter tracts and 20 cell types across 9 microstructural measures, based on Spearman’s Rho and associated p-values corrected for multiple comparisons across cell types (Benjamini–Hochberg FDR; q<0.05).

The association between the 20 cell types and enrichment estimates across 9 age-linked microstructural measures in 48 whole brain white matter tracts is illustrated in Figure 5A (Detailed results are presented in Supplementary Table 4). Readers should note that the results of these analyses represent cellular associates of white matter tracts in aging through nomination of gene sets spatially related to cell types. Critically, while these data may suggest the presence of relationships linking the integrity of white matter tract to specific cell types, they should not be taken to imply the presence of cells located within these tracts. Enrichment estimates for oligodendrocytes, a chief cell component present in cerebral white matter, showed significant association with age-related microstructural measures across whole brain tracts, including FA (Spearman’s *r_s_*=-0.55, p_FDR_=6.328e-4) and diffusion tensor eignenvalues L1 (Spearman’s *r_s_*=0.40 p_FDR_=0.005) and L2 (Spearman’s *r_s_*_=_0.52, p_FDR_=0.003; Figure 5B). While oligodendrocytes represent a primary cell target of interest in white matter development and aging, the supporting role of other glial cells should not be overlooked. As shown in Figure 6, enrichment estimates for glial cells such as astrocytes and microglia were positively associated with NODDI diffusion measures, such as ISOVF (astrocytes: Spearman’s *r_s_*_=_0.45, p_FDR_=0.027; microglia: Spearman’s *r_s_*_=_0.44, p_FDR_=0.027), and OD (microglia: Spearman’s *r_s_*_=_0.43, p_FDR_=0.045). The relationship between astrocyte enrichment and ICVF displayed a similar, but trend-level, pattern that did not survive multiple comparison correction (Spearman’s *r_s_*_=_0.39, p_FDR_=0.067). Endothelial cell enrichment was found to be significantly positively associated with ICVF (Spearman’s *r_s_*_=_0.44, p_FDR_=0.039). Though historically overlooked (but see pioneering studies examining astrocyte modulation on synapse signaling) (79, 80), recent work has demonstrated the regulatory role of these non-neuronal support cells in the health of white matter microstructure. Growing evidence supports the notion of an astrocyte-microglia axis, where these cell types engage in cross-talk to maintain homeostasis and engage in tandem in the presence of pathology, where coordinated loss in supportive functions and morphological changes occur in favor of optimizing survival (81–83). The response activation of microglia and astrocytes also features as a prominent hallmark in aging and progression of Alzheimer’s disease, for instance, their crucial role in the mechanisms underpinning neuroinflammatory processes, which in turn, impact functional integrity of surrounding brain cells (68, 84).

**Figure 5.**
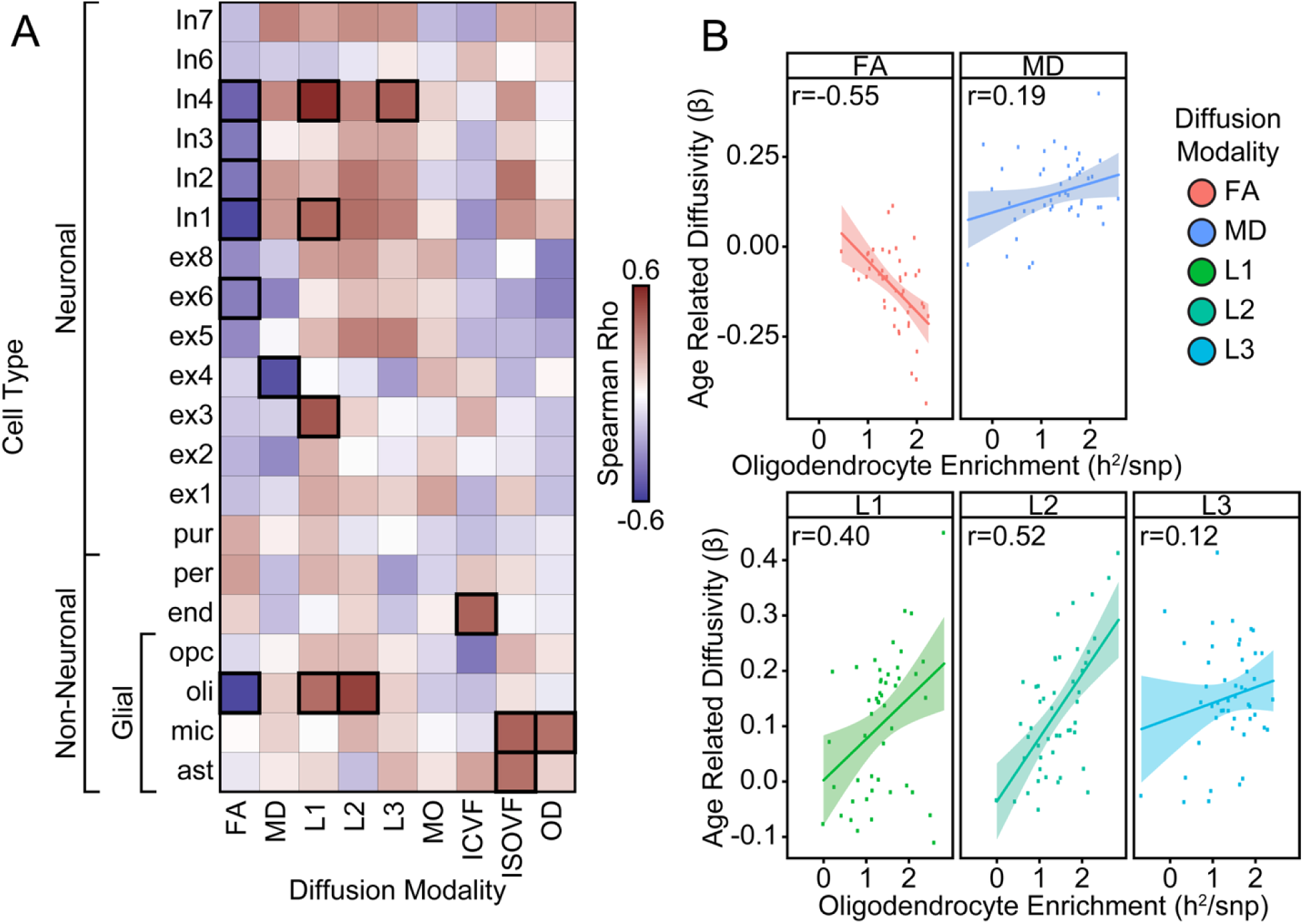
Cell type distributions track with age-related associations across diffusion measures. Cell type enrichment estimates were calculated based on compiling differentially expressed marker genes across cell subtypes identified by Lake et al (78), using LDSC (linkage disequilibrium score regression) partitioned heritability. Twenty cell types were rank ordered by spatial correlation to whole brain white matter tracts across diffusivity measures. (A) Table cells reflect correlations between age-regressed estimates and cell type enrichment, calculated by Spearman’s Rho. Warm red colors indicate positive correlation, cool blue colors indicate negative correlation, and significant correlations that survived FDR correction (p<0.05) are highlighted by bolded cells. (B) Scatter plots colored by the 20 respective diffusivity measure depict the positive/negative relationship between age-regressed diffusion estimates and oligodendrocyte enrichment (FA=fractional anisotropy, MD=mean diffusivity, L1, L2, L3 representing individual eigenvalues λ1, λ2, λ3 relative to the diffusion tensor). Oligodendrocyte enrichment was significantly negatively correlated with FA, and inversely correlated for the other diffusivity metrics, such as MD and L1-3.

**Figure 6.**
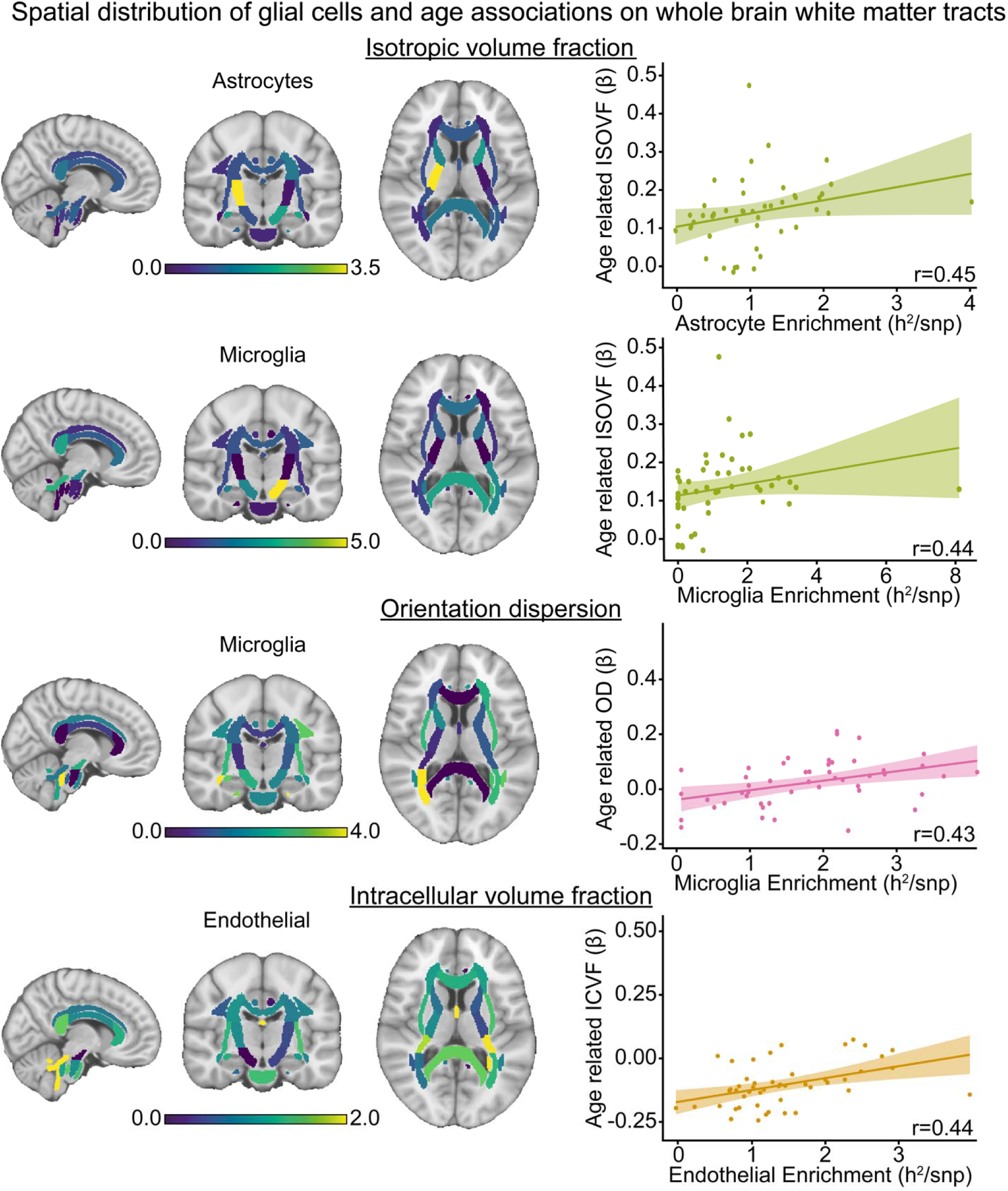
Glial cell distribution track with age-related NODDI diffusion measures. Spatial distribution of glial cell associations with white matter (left panel) and corresponding age associations (right panel) across whole brain white matter tracts for NODDI diffusion estimates (i.e., ISOVF=isotropic volume fraction; OD=orientation dispersion; ICVF=intracellular volume fraction). Scatter plots colored by the respective diffusivity measure show the positive relationship between age regressed NODDI diffusion estimates (b) and glial cell enrichment, such as astrocytes and microglia. Correlations were estimated by Spearman’s Rho.

Of note, the observed relationships are not exclusive to non-neuronal glial cells. Here, we highlight associations that were consistently identified between several age-linked diffusion measures and enrichment in neuronal cells. Namely, these include inhibitory interneuron cells such as In1 and In4. Both In1 and In4 interneuron cell types were found to be significantly negatively associated with age-related FA (In1: Spearman’s *r_s_*_=_0.43, p_FDR_=6.3e-4; In4: Spearman’s *r_s_*_=_-0.46, p_FDR_=0.007) and positively associated in diffusion tensor eigenvalues L1 (In1: Spearman’s *r_s_*_=_0.43, p_FDR_=0.002; In4: Spearman’s *r_s_*=0.57, p_FDR_=5.7e-4) and L3 (In4: Spearman’s *r_s_*_=_0.44, p_FDR_=0.039) respectively. Sparse but significant relationships were also observed for excitatory cells Ex4 and Ex6 with age-associated MD (Spearman’s *r_s_*_=_-0.52, p_FDR_=0.004) and FA (Spearman’s *r_s_*_=_-0.37, p_FDR_=0.032). In1 comprises a large group of inhibitory interneurons that are made up of multiple subtypes (i.e., In1a, In1b, In1c) marked by high expression of genes such as VIP, RELN, cholecystokinin (CCK), and cannabinoid receptor 1 (CNR1). RELN also makes up part of inhibitory cell type In4 (comprising subtypes In4a and In4b). It should be noted that based on chromatin accessibility signatures associated with transcriptional subpopulations, the In4 cluster constructed in the original dataset by Lake and colleagues (78) were not consistently distinguishable from In1,2,3 clusters, suggesting a lack of differentially accessible sites sufficient to reliably resolve the In4 cell partition versus the rest of the In1-3 subgroups. As a whole, these data demonstrate the preferential association of cell types that explain a substantial portion of the heritable variation across a range of microstructural diffusion MRI measures that track age-associated changes in white matter composition.

## Discussion

Recent progress in human neuroscience has provided the opportunity to integrate *in vivo* estimates of brain function and anatomy with *ex vivo* measures of molecular and cellular processes. Here, leveraging analyses of heritability in a large aging cohort, we demonstrate that a substantial portion of population-level variability in diffusion-based measures of tract microstructure is under the influence of genetic factors. Across microstructural measures, while individual tracts display variation in their heritability, genetic similarity is greater within, than between, broad white matter tract classes defined based on their patterns of connectivity. Late-life aging shows preferential reductions in white matter microstructure within association tracts, relative to other tract classes. Cell type analyses implicate oligodendrocytes and other glial cells as possessing a preferential, although not specific, relationship with age-related changes in microstructure across whole brain tracts. These enrichment-to-age associations suggest that white matter tracts whose variability is most strongly influenced by genes expressed in a given cell type may also be the tracts most vulnerable to age-related decline. For example, the observation that oligodendrocyte enrichment tracks with age-related reductions in tract integrity implies that cellular processes governing myelination may contribute disproportionately to late-life white matter deterioration. This framework provides a biologically grounded link between inter-individual variability in tract integrity and the cellular substrates of white matter aging.

The structural wiring properties of the human brain are under genetic control. Consistent with prior work (10, 11, 46, 85), our findings demonstrated strong genetic constraints on white matter microstructural metrics such as ICVF and FA (median h^2^ ∼ 0.25-0.35) (10, 86). In a large population-based sample (N=29,942), our data reveal diffusion-based measures of white matter microstructure possess moderate to high heritability. White matter pathways serve as an anatomical backbone, supporting the coordinated communication of information across spatially distributed functional systems (87–89). Convergent evidence in human and non-human animal models and across spatiotemporal scales has revealed a strong correspondence between brain structure and function, from local circuits and neural populations (90) through the fiber bundles that connect large-scale brain networks (91, 89, 92). However, while the organizational properties of white matter connections in the human brain have been mapped across levels of analyses, both in *in vivo* and in postmortem tissue samples, the relations linking genetic and cellular properties of white matter organization with microstructural measures derived from non-invasive diffusion MRI remains to be established.

Suggesting the presence of common genetic influences underlying white matter tracts with shared patterns of structural connectivity, our data indicate that estimates of genetic similarity are significantly higher within, than between, major, commonly reported tract classes (complex, associative, commissural, and projection; Figures 2, 3). This property of brain organization was broadly evident across measures of white matter microstructure derived by classical DTI (e.g. FA, MD) and NODDI models (Supplementary Table 3). Taken together, these data provide evidence for shared genetic factors underlying the organizational properties of large-scale fiber bundles. Suggesting potential developmental and evolutionary origins of the shared genetic bases, fiber tracts of the human brain appear to be organized in a lamellar pattern (93, 94), akin to that of the laminar origin of cortico-cortical connections observed in nonhuman primates (95). Connectomic studies in the human and non-human mammalian brain have revealed that cortical regions that are structurally connected by axon projections are more likely to share similar features in cytoarchitecture, such as neuronal density, laminar differentiation, and profiles of gene expression (21–25). Moreover, cytoarchitectural similarities among connected brain regions likely reflects greater levels of coordinated cortical maturation (26, 27).

Developmental and lifespan trajectories of white matter tract maturation and maintenance follow complex spatial and temporal patterns, where white matter changes are observed to be non-uniform across the brain and follow a protracted course of maturation (40, 41, 71). During ontogenesis, for example, the rate of myelination is more rapid in motor root than sensory system fibers (96, 97). Temporally offset from the maturation of unimodal systems, the exponential growth of association fiber pathways is evident well into adolescence extending into early adulthood (41, 98, 99). Work in aging populations has lent support to the “retrogenesis” hypothesis, which postulates that tracts that mature later such as association tracts, tend to be more susceptible to earlier deterioration in late life (24, 42–44, 100). One suggested theory for this pattern is that the pathways latest to develop are more thinly myelinated and that the mature oligodendrocytes responsible for their myelination are more vulnerable to accumulating deleterious effects due to their elevated metabolic activity (42).

Consistent with this literature, our analyses indicate that association fiber tracts display greater age-related vulnerability, evidenced by marked changes across measures of white matter microstructure, relative to other tract classes (Figure 4 and Supplementary Figure 2). For instance, tracts such as the fornix, a key output tract of the hippocampal formation, exhibited marked susceptibility to aging effects in the present data. The fornix serves as a crucial structural phenotype for brain age predictions across diffusion MRI approaches (101), and is implicated in cognitive faculties such as episodic memory and recall (102, 103). Considering these age-related effects can be observed within white matter tract classes defined by broad connectivity profiles, as well as our findings demonstrating genetic similarity within these tract classes, these data suggest that common genetic factors underlying white matter tracts capture shared patterns of structural brain connectivity that has been shown to track with age-related vulnerability.

An improved understanding of the biological principles governing dynamic alterations in brain white matter microstructural properties across the lifespan is critical for understanding the process of age-related changes in cognitive functioning and for the eventual development of associated precision medicine approaches. Changes in brain white matter microstructural properties are dynamic across the lifespan and likely do not occur in a homogeneous fashion. Overall, older age was associated with significant changes across white matter microstructural indices such as FA and MD, as well as in NODDI indices (Supplementary Data S11), broadly indicating poorer white matter health. Bridging *in vivo* estimates of white matter microstructure with genetics and cell typing data, we provide evidence suggesting the genetic associates of glial cells account for a substantial portion of the heritable variation in a range of *in vivo* white matter microstructural measures. Longstanding histological evidence harking back to staining visualizations by pioneers Hortega and Penfield have revealed that myelin, formed by oligodendrocytes enriched in brain white matter, sheath axons to promote conduction and provide metabolic support to neurons, influencing communication between brain regions and neural circuit function (104). The advent of non-invasive diffusion MRI and accompanying computational models provided the opportunity to estimate measures sensitive to features of white matter tissue such as axonal configuration, packing density, and myelin density (105, 106). Our analyses build on these advances to provide molecular genetic support for a relationship between glial cells and alterations in white matter tract microstructure in the aging human brain. Notably, polymorphisms linked to gene markers of oligodendrocytes accounted for an enriched proportion of heritable variance underlying age-related white matter microstructural changes, as indexed by DTI measures such as FA, L1 and L2 (Figure 5).

Critically, our results also highlight the implication of glial cells such as astrocytes and microglia in the aging of brain white matter. Multi-compartment models of diffusion such as NODDI can potentially provide more specific markers of brain tissue microstructure than traditional DTI (107). In particular, variance explained by SNPs linked to genetic markers of glial cell types significantly tracked with age-linked changes in NODDI measures such as ISOVF and OD (Figure 6). This is in support of hypotheses theorizing the role of glia in cell homeostasis and inflammation modulatory responses in maintenance of the health of white matter microstructure (108, 109). These findings are consistent with recent work establishing associations between NODDI measures such as ISOVF and ICVF with markers of inflammation with genes enriched for immunity and neurotransmitter receptors (110, 111). Converging evidence in rodents examining finer-grained spatiotemporal transcriptomics, has identified brain-wide aging signatures in glial cells exhibiting accelerated changes in white matter relative to cortical regions (112). Aging induces spatially dependent cell-state changes in non-neuronal cells, underscoring white matter as a ‘hotspot’ for aging-related glial and immune cell activation (30). These data are in line with evidence for associations linking Alzheimer’s disease with impaired microglia functioning and neuroinflammation (68, 84), suggesting a role for both myelin formation and cellular processes that support neuron health in the maintenance of tract fibers in late life. Taken together, these data suggest that the common genetic polymorphisms that influence age-linked changes in the microstructural properties of brain white matter do so in a cell-preferential manner.

Of note, the observed cellular associates of white matter changes in late life were not specific to glial cells. We also observed significant relationships between age-related white matter microstructural measures and neuronal cells, particularly for inhibitory cell types such as In1, In4, and excitatory cell types such as Ex4 and Ex8. In animal models, axonal myelination of inhibitory and excitatory neurons is regulated by activity and experience, involving alterations in axonal structure and myelin. Such activity-dependent plasticity suggests that age-related shifts in neuronal firing may adaptively remodel white matter across the lifespan (113). For instance, the firing and synaptic properties of VIP cells, a class of In1 interneuron cells, are significantly altered in senescent mice (114, 115). Furthermore, although speculative, these non-glial cell associations with white matter may emerge through secondary biological processes, or downstream cascades that indirectly influence tract microstructure. These could include, for example, the functioning of local micro-circuits that support the health of pyramidal cells and maintenance of associated axonal projections. Additionally, a sparse but growing body of evidence suggests the presence of interstitial neurons within subcortical white matter, where this subset of cortical neurons possesses similar morphological and molecular diversity to overlying cortical neurons, expressing glutamatergic and GABAergic markers (20, 116–118). These cells have been reported to be integrated in cortical circuitry and be able to actively coordinate interareal connectivity (20). Although speculative, the present data may, in part, reflect this aspect of brain organization.

The present analyses provide a novel first step in bridging *in vivo* imaging findings with cellular processes for the study of structural white matter changes in the aging brain. However, the present study should be interpreted in light of several limitations. First, we utilized nominated cell type gene marker sets based on predefined statistical clustering (78). Although we are unable to conclude that genes within each imputed cell type group directly influence their respective functions, similar correlation-based nomination approaches have been reported to correspond well with *a priori* defined gene groups (21, 119). In addition, the cell types analyzed were derived from a limited set of brain regions, namely, prefrontal and visual cortices, and do not provide a complete characterization of the spatial variation of cell type profiles throughout the human brain. Furthermore, our present analyses are based on assessing multiple white matter microstructural measures of large fiber bundles ranging from conventional DTI to more complex parameters derived from NODDI. While providing in vivo estimates of tissue microstructure, this approach may not fully capture topological or network connectivity properties that alternate methods such as tractography may offer. Since tractography allows for the delineation of white matter connections and estimation of connectivity strengths, interindividual variability in topological features of the white matter connectome may be influenced by partly distinct genetic variants from those related to white matter microstructure (18). Future work could consider integrating across methodologies to provide complementary insights. For example, distinct genetic and cellular influences on axon outgrowth during the development of long-range connections may be detectable through estimates of connection strength as measured through tractography, relative to indices of microstructural integrity. An alternative but complementary approach could be to apply cell type enrichment directly to GWAS summary statistics for age associated brain phenotypes, as in work by Smith and colleagues (120), which identified genetic variants influencing brain aging. Finally, it is crucial to underscore that although we classified white matter tracts into canonical anatomical categories, this is a simplified taxonomy. Thus, although our classification may provide a high-level framework for investigating predominant connectivity patterns, it does so by grouping highly heterogeneous white matter pathways. This reflects a broader issue in *in vivo* neuroimaging: macro-scale atlases often average over fine-grained anatomical complexity and therefore, interpretations of these results stratified by tract-class should be viewed with this caveat in mind.

Inherited genetic factors influence the formation and maintenance of white matter tracts across the lifespan. In a well-powered community cohort, we first validated the genetic landscape of *in vivo* microstructure across white matter tracts via SNP heritability and established the presence of shared genetic factors underlying broad categories of major fiber bundles. We then localized and quantified age-related differences in white matter tracts and confirmed the presence of age-linked vulnerability in specific groups of tracts, particularly association tracts. Through leveraging multi-scale data from neuroimaging, genetics, and transcriptomics, we further demonstrate that age-related changes across a range of diffusion-based estimates of microstructure are preferentially linked with gene markers of cell types, such as glial cells. Collectively, these findings demonstrate the genetic and cellular underpinnings of brain structural wiring properties in late life and offer a neurobiological basis for further planned exploration of risk factors and molecular mechanisms of brain aging.

## Methods

### UKB sample

This study was conducted under UK Biobank application 25163.The UK Biobank received ethical approval from the National Research Ethics Service Committee North West-Haydock (reference 11/NW/0382), and all procedures were performed according to the World Medical Association guidelines. All enrolled participants provided written informed consent. In line with the goals of the current study, we primarily utilized the diffusion MRI (dMRI) data (UK Biobank data field: 20250, first imaging visit) released in February 2020, with accompanying genome-wide genotyping array data. All the following analyses were conducted in accordance with the guidelines of the Yale University Institutional Review Board (IRB).

### UKB neuroimaging

Structural MRI data from the UK Biobank were acquired and analyzed using the standard UKB brain imaging protocol (48). Briefly, brain data were acquired on a single standard Siemens Skyra 3T scanner with a standard Siemens 32-channel RF receiver head coil, with the imaging matrix angled down by 16 from the AC-PC line. The T1-weighted volumes were acquired in the sagittal plane using a three-dimensional magnetization-prepared rapid gradient-echo sequence at a 1×1×1mm resolution, with a 208×256×256 field of view. The dMRI protocol employed a spin-echo echo-planar imaging sequence with 10 T_2_-weighted (b»0s mm^2^) baseline, 50 b=1,000s mm^-2^, and 50 b=2,000s mm^-2^ diffusion-weighted volumes acquired with 100 distinct diffusion-encoding directions and multi-slice acceleration factor 3. The field of view was 104×104mm, imaging matrix 52×52, 72 slices with slice thickness 2mm, giving 2mm isotropic voxels. White matter tracts were derived from tract-based spatial statistics (TBSS) analyses (121), where a standard-space white matter skeleton was mapped to each subject using a high-dimensional warp, before 48 individual regions of interests were delineated based on the skeleton with standard-space masks from the JHU ICBM-DTI-81 white matter atlas (122). Across these 48 white matter tracts, nine diffusion metrics (FA, MD, L1, L2, L3, MO, ICVF, ISOVF, and OD) were calculated by the UK Biobank imaging pipeline and made available as 432 unique imaging-derived phenotypes (IDPs) of white matter microstructure (48). Additionally, white matter tracts were broadly classified into four main groups: complex, associative fibers, commissural fibers and projections (refer to Supplementary Table 2 for a complete classification list).

Participants who reported diagnoses of neurological disorders such as dementia/Alzheimer’s disease, Parkinson’s disease, stroke diagnoses or any other chronic/degenerative neurological or demyelinating diseases such as multiple sclerosis were excluded. Participants were filtered for missing and/or quality data and excluded based on outliers for imaging variables such as number of dMRI slice outliers and white matter lesions, where individuals with ±3 SD from the mean were removed. Extreme outlier participants that had average FA measuring ±4 SD from the mean were also excluded from further analyses.

### UKB genetics and SNP-based heritability

To ensure a high degree of genetic homogeneity, UK Biobank genotype data was filtered to include only White British participants with imaging data passing the quality control thresholds described above. Amongst individuals with available dMRI and genotype data, we excluded participants with a mismatch of their self-reported and genetically inferred sex, with putative sex chromosome aneuploidies, or who were outliers according to heterozygosity (±3 SD). The final inclusion sample for subsequent analyses consisted of 29,942 participants (filtered from 38,935), with a mean age of 63.1 years (age range=45-81 years) and was 53.3% female.

Plink v2.0 was used to remove samples with genotype missingness >0.10, SNPs with minor allele frequency <0.05, Hardy–Weinberg equilibrium p < 1×10^−6^, and call rate <0.02, resulting in 337,501 autosomal variants (123). GCTA software (v.1.93.0) was used to calculate a genetic relatedness matrix to remove individuals with cryptic relatedness more than 0.025, as SNP-based heritability analysis is especially sensitive to participants with higher levels of relatedness (124). Ten genetic principal components were then calculated for use as covariates in subsequent genetic and heritability analyses.

Genome-wide association analyses were conducted using the linear regression form of GCTA’s *fastGWA* utility (105). Age, sex, height, weight, combined normed gray/white matter volume (head size normalized), T1 inverse SNR (signal to noise ratio), number of dMRI outlier slices, T2-weighted FLAIR white matter lesion volume, head coordinates (*X, Y, Z*), UK Biobank assessment center, and the 10 genetic principal components were included as covariates. Quantitative variables were *z*-transformed. SNP heritability of 432 white matter microstructural tract measures were estimated with genotyped data using GCTA-REML and replicated in parallel using LDSC regression (125) using the same quantitative and categorical covariates above. LDSC based estimates of white matter microstructural heritability were conducted on 1,158,800 HapMap3 variants that overlapped with variants in the UKB GWAS summary statistics, using precomputed LD scores from 1000 Genomes European data (i.e., “*eur_w_ld_chr*”) (126).

### Genetic correlation between white matter tract phenotypes

Genetic correlation analyses were performed using the GCTA bivariate genomic-relatedness-based restricted maximum likelihood (GREML) utility (127, 128), which captures SNP-based estimation of genetic variance explained by phenotypic correlations between traits. Bivariate genetic correlation scores (r_g_) were calculated between 48 pairs of white matter tracts for each diffusion measure, repeated across the 9 measures. For all analyses, we included the same categorical and quantitative covariates as in the GWAS and heritability analyses. To examine if genetic correlation was significantly greater for pairs of white matter tracts classified within the same tract class compared to pairs that belonged to separate distinct classes, within- and between-group differences for genetic correlation estimates (r_g_) were tested using paired sample t-tests.

### Regression linking aging to structural neuroimaging phenotypes

Regression analyses were conducted independently across the 9 diffusion measures for 48 white matter tracts. Quantitative variables were *z*-transformed. The effect of age on the respective white matter diffusion measures were estimated, covarying for height, weight, sex, age, age^2^, age*sex, combined gray/white matter volume (head size normalized), T1 inverse SNR (signal to noise ratio), number of dMRI outlier slices, diastolic and systolic blood pressure, head position coordinates (*X, Y, Z*) of brain in the scanner (center mass of brain mask), and UKB imaging acquisition center. Tests of the difference between age association magnitudes between tract classes were conducted on the linear component of tract measures, implemented using paired sample t-tests.

### Single-cell transcriptional enrichment analyses

To infer associations of cell type abundance, we utilized single-nucleus droplet-based sequencing (snDrop-seq) data from Lake and colleagues (78), noting that similar approaches with imputed cell-type distributions have previously been shown to align with macro-scale cortical organization (129). Specifically, a select pool of 8,066 significantly differentially expressed genes between all snDrop-seq clusters across frontal and visual cortices identified as significant enriched cell type gene markers distinguished by their spatial orientation (made publicly available in Supplementary Table S3) were used for the following analyses (78). To reduce collinearity, the 20 superordinate cell identities as originally defined were applied in categorizing transcriptionally similar cell types.

The *biomaRt* package in R (130) was used to map SNP locations to genes and to identify each gene’s transcription start and stop sites (±10,000 base pairs) according to the AHBA Entrez IDs (i.e. GRCh37-hg19 genome assembly build). Entrez IDs were matched and de-duplicated, genes without Entrez IDs were removed. A small subset of genes did not have analyzable variants using these criteria and did not contribute to results.

Partitioned heritability analyses were performed for the 432 white matter microstructural tract phenotypes. This was done by examining SNPs occurring in spatial proximity to the coding regions of each gene bin corresponding to cell type identities. Intragenic (±10,000 base pairs) and eQTL SNPs associated with the gene bins were used for partitioned heritability analyses.

The LDSC “*make_annot.py*” tool was used to create annotation files, filtering on HapMap3 variants and using Phase 1 genetic data from the 1000 Genomes Project to estimate LD. Partitioned heritability of the derived SNP sets was estimated in conjunction with a full baseline model (i.e., “*1000G_Phase1_baseline_ldscores*”), using precomputed allele frequencies (i.e., *1000G_frq*) and weights (i.e., “*weights_hm3_no_hla*”). These analyses provide a measure of genetic variance explained by the SNP bins corresponding to the 20 cell type identities, conditioned on the baseline annotation model to prevent non-specific genetic signals from inflating stratified heritability estimates. Enrichment values calculated in partitioned heritability analyses were defined as:

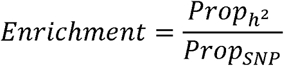

where 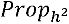 is the phenotypic variance explained by SNPs associated with the individual cell type / total SNP heritability, and *Prop_SNP_* is the number of SNPs associated with the individual cell type / total number of SNPs used in the analysis. Thus, the enrichment values provide an estimate of whether a specific cell type has a disproportionate contribution to overall SNP heritability. Since heritability is a variance explained measure and in theory non-negative, heritability enrichment estimates < 0 were set to zero.

### Identification of cell types spatially correlated to age-associated white matter microstructure

We investigated whether enriched gene markers nominated by single-cell data from Lake et al (78) are spatially linked with measures of whole brain white matter microstructure. Specifically, the spatial relationship between enrichment estimates of the 20 cell types and 432 age-associated white matter tract microstructural phenotypes was quantified using correlation (*Spearman’s* Rho). P-values were corrected for multiple comparisons (Benjamini–Hochberg false-discovery rate; q <0.05) across cell types.

## Acknowledgements

This work was supported by the National Institute of Mental Health (Grants R01MH120080 and R01MH123245 to AJH), Tan Kah Kee Foundation, Singapore (Tan Ean Kiam Postgraduate Scholarship Award to RC). The data used in this work were obtained from UK Biobank (Data Application 25163). We are grateful to UK Biobank for making the data available and to all UK Biobank study participants, who generously donated their time to make this resource possible. This work also used publicly available data from the CommonMind Consortium. We thank Tian Ge, Sidhant Chopra, and Ashlea Segal for their feedback on analyses and prior versions of the project. Analyses were made possible by support from high-performance computing facilities provided through the Yale Center for Research Computing.

