## Supplementary Information for "The cellular associates of late life changes in white matter microstructure"

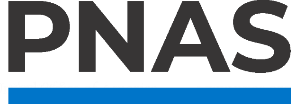

**Supporting Information for**

The cellular associates of late life changes in white matter microstructure

**Authors:** Rowena Chin^1,2^, Xi-Han Zhang^1,2^, Kevin M. Anderson^1^, Anastasia Yendiki^3^, Avram J. Holmes^4^

**Affiliations:**

^1^ Department of Psychology, Yale University, New Haven, CT, USA

^2^ Wu Tsai Institute, Yale University, New Haven, CT, USA

^3^ Athinoula A. Martinos Center for Biomedical Imaging, Department of Radiology, Massachusetts General Hospital/Harvard Medical School, Charlestown, MA, USA

^4^ Department of Psychiatry, Brain Health Institute, Rutgers University, Piscataway, NJ, USA

**Correspondence:** Rowena Chin, Yale University, Department of Psychology, 100 College St, New Haven, CT 06520,. Avram J. Holmes, Rutgers University, Department of Psychiatry, Brain Health Institute, 119 Staged Research Building, 661 Hoes Lane West, Piscataway, NJ 08854,.

**This PDF file includes:**

Supporting Text

Figures S1 to S11

Tables S1 to S4

SI References

Supporting Text

**Extended description of diffusion metrics**

FA and MD respectively capture the directional coherence and magnitude of water molecule diffusion in tissue (1). Water molecules tend to diffuse with greater directional coherence and lower magnitude when constrained by tightly packed fibers such as well-myelinated axons and by cell membranes, microtubules, and other cellular structures. Based on evidence from post-mortem and tract-tracing studies, individual differences in measures of diffusivity are theorized to reflect meaningful differences in underlying brain white matter microstructure (2, 3). Other related measures of tract-averaged L1, L2, L3, and MO are also parameters of interest to brain aging; for example, L1, L2, L3 are the three main tensor eigenvalues, from which FA and MD are computed (1), and MO describes the type of anisotropy as a continuous measure reflecting tensor shape differences that range from planar (e.g. in regions of crossing fibers) to linear (e.g. in regions where one fiber population predominates) (4, 5). Unlike DTI, which fits a Gaussian distribution to the diffusion MRI signals, NODDI fits a multi-compartment model that decomposes the diffusion MRI signal into: free water diffusion (e.g. in CSF), restricted (intra-axonal) diffusion and hindered (extra-axonal) diffusion (6). The resultant indices are ICVF (the contribution of intra-axonal diffusion, i.e., a measure of neurite density), ISOVF (the contribution of free water diffusion), and neurite OD (the degree of dispersion or fanning in neurite orientation) (6). While DTI measures such as FA are not able to discriminate between different biological changes that may lead to a change in anisotropy (7), the NODDI measures may offer greater specificity in characterizing white matter aging.

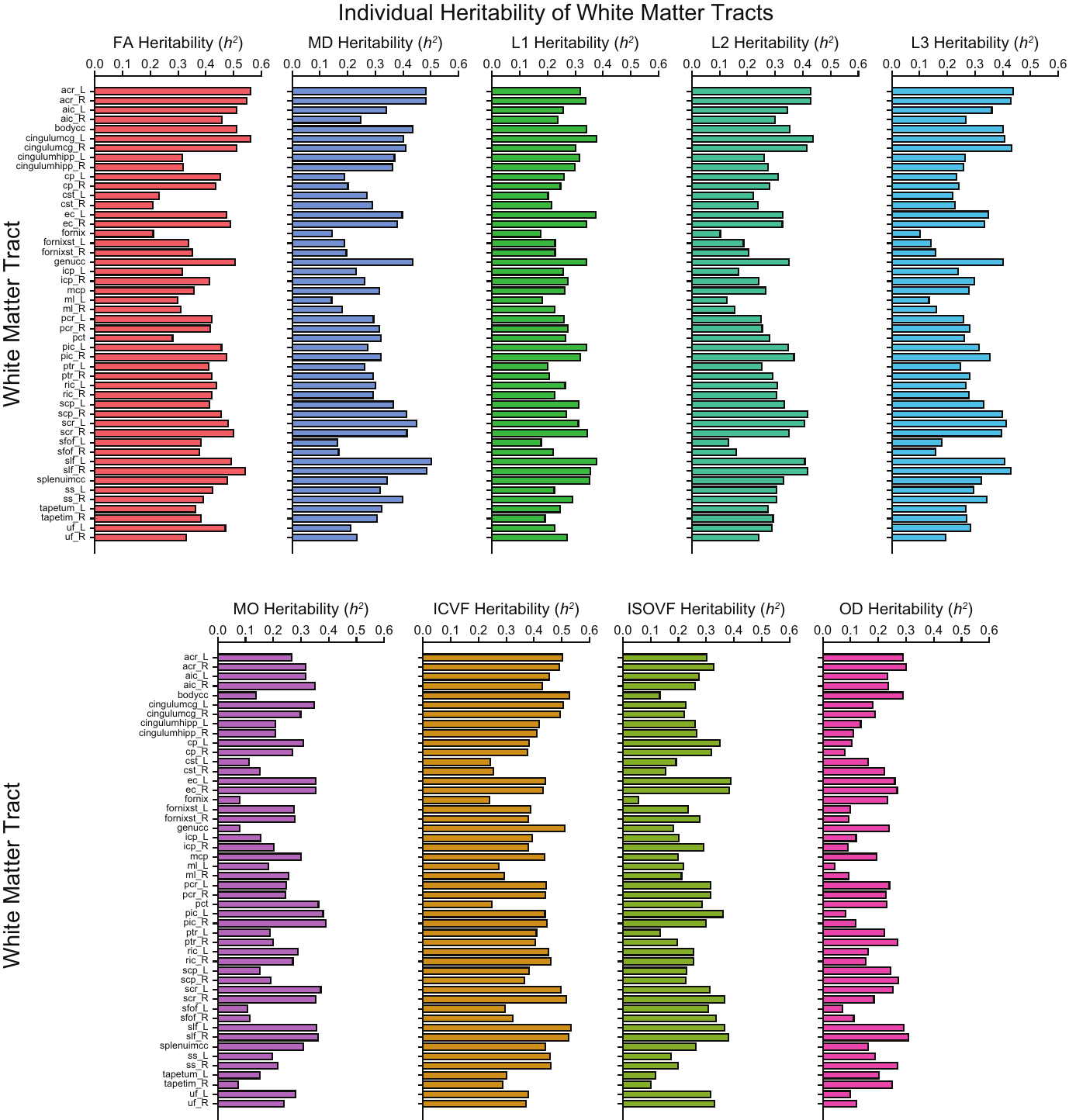

**Fig. S1.** Heritability estimates (h^2^) across 48 white matter tracts labeled based on the JHU ICBM-DTI-81 White Matter Atlas for 9 individual diffusion MRI modalities (totaling 432 diffusivity-based tract measures). Individual boxplots are colored according to each diffusivity modality.

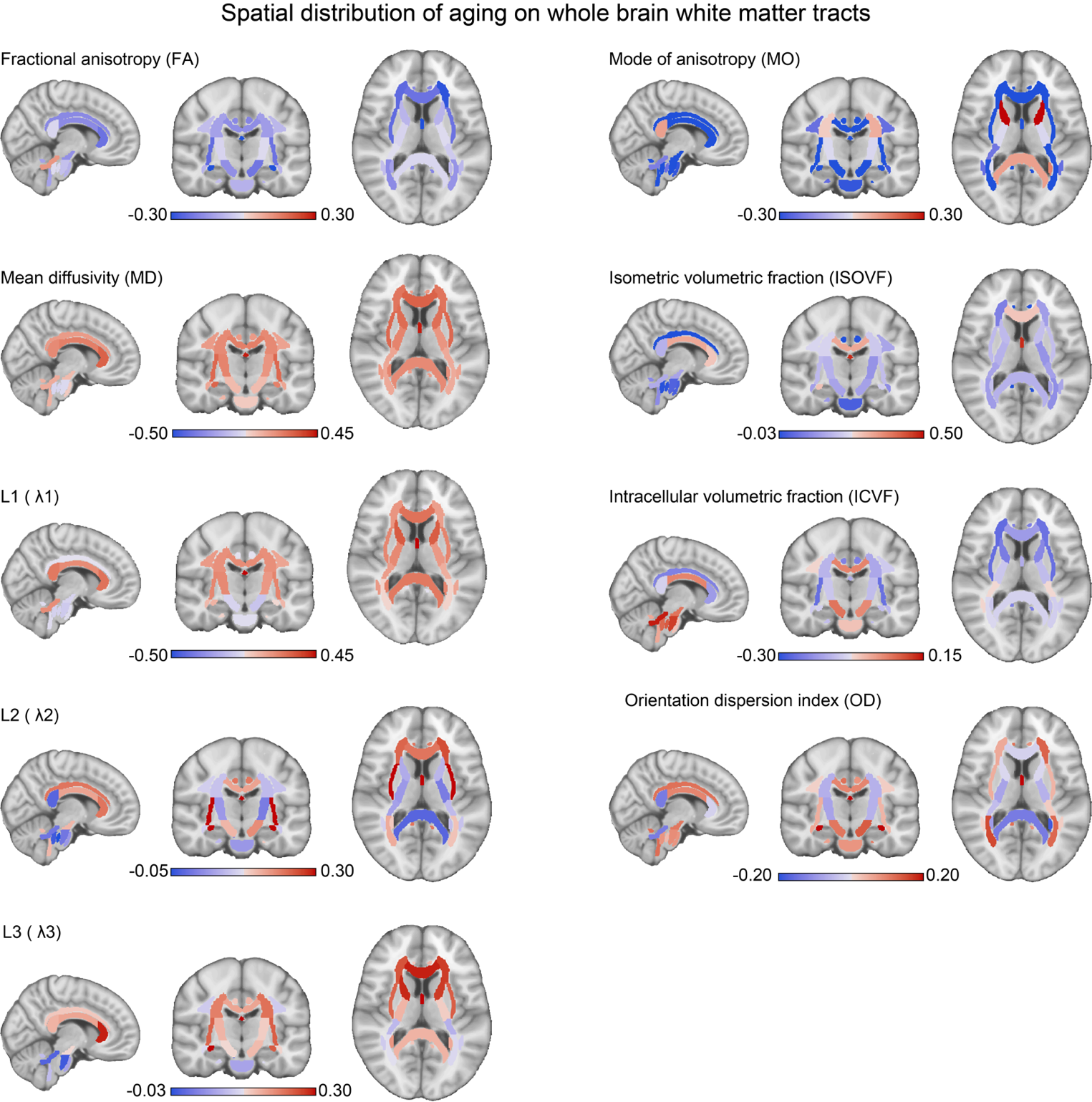

Fig. S2. Spatial distribution of aging effects on whole brain white matter tracts across 9 diffusion MRI measures. Positive age associations were marked in red, while negative age associations were marked in blue.

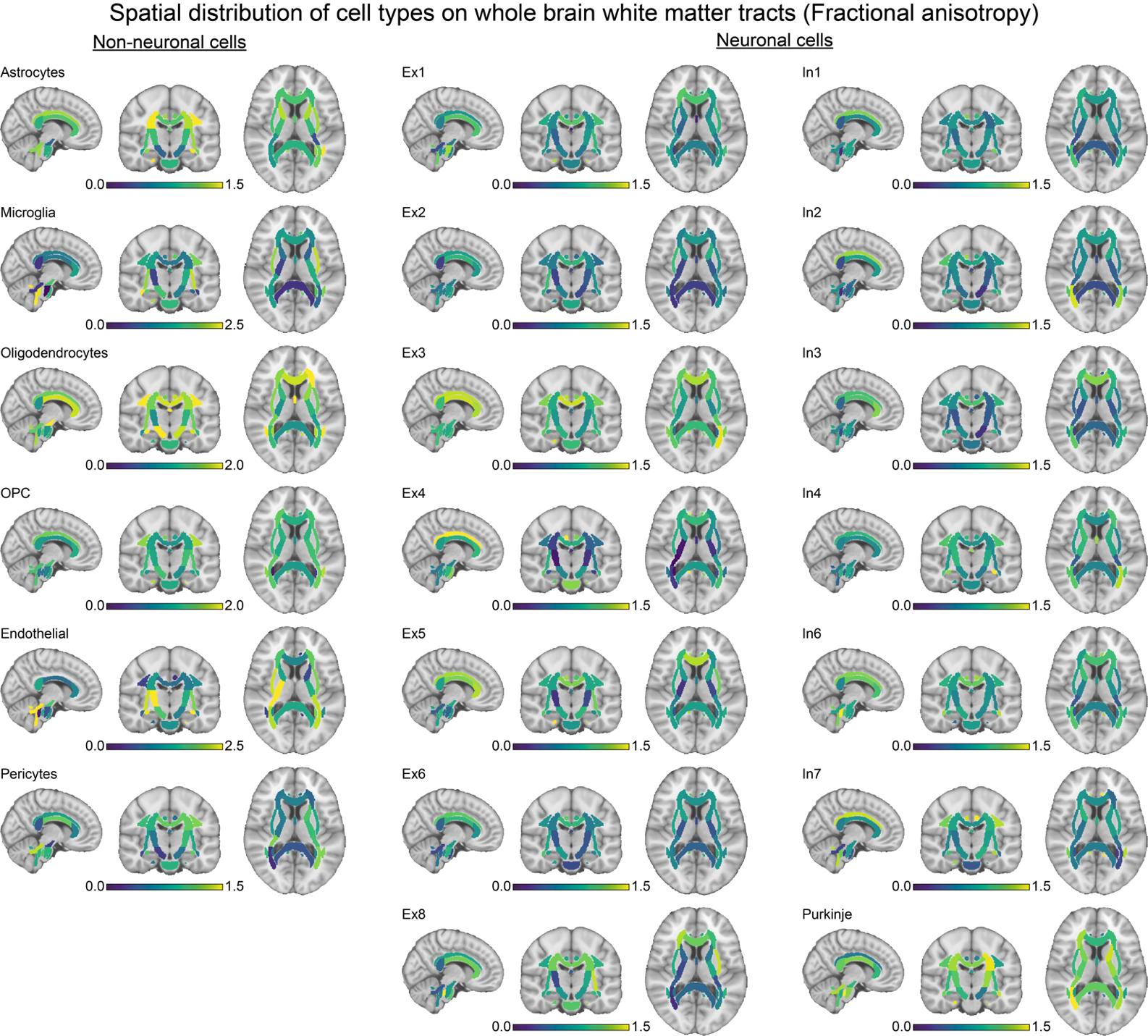

**Fig. S3.** Spatial distribution of cell type enrichment on whole brain white matter tracts as measured by fractional anisotropy (FA). Individual cell type enrichment distributions across twenty cell subtypes are sectioned into neuronal (right panel) and non-neuronal groups (left panel) and displayed accordingly. Dark blue color indicates low enrichment, while bright yellow indicates high enrichment.

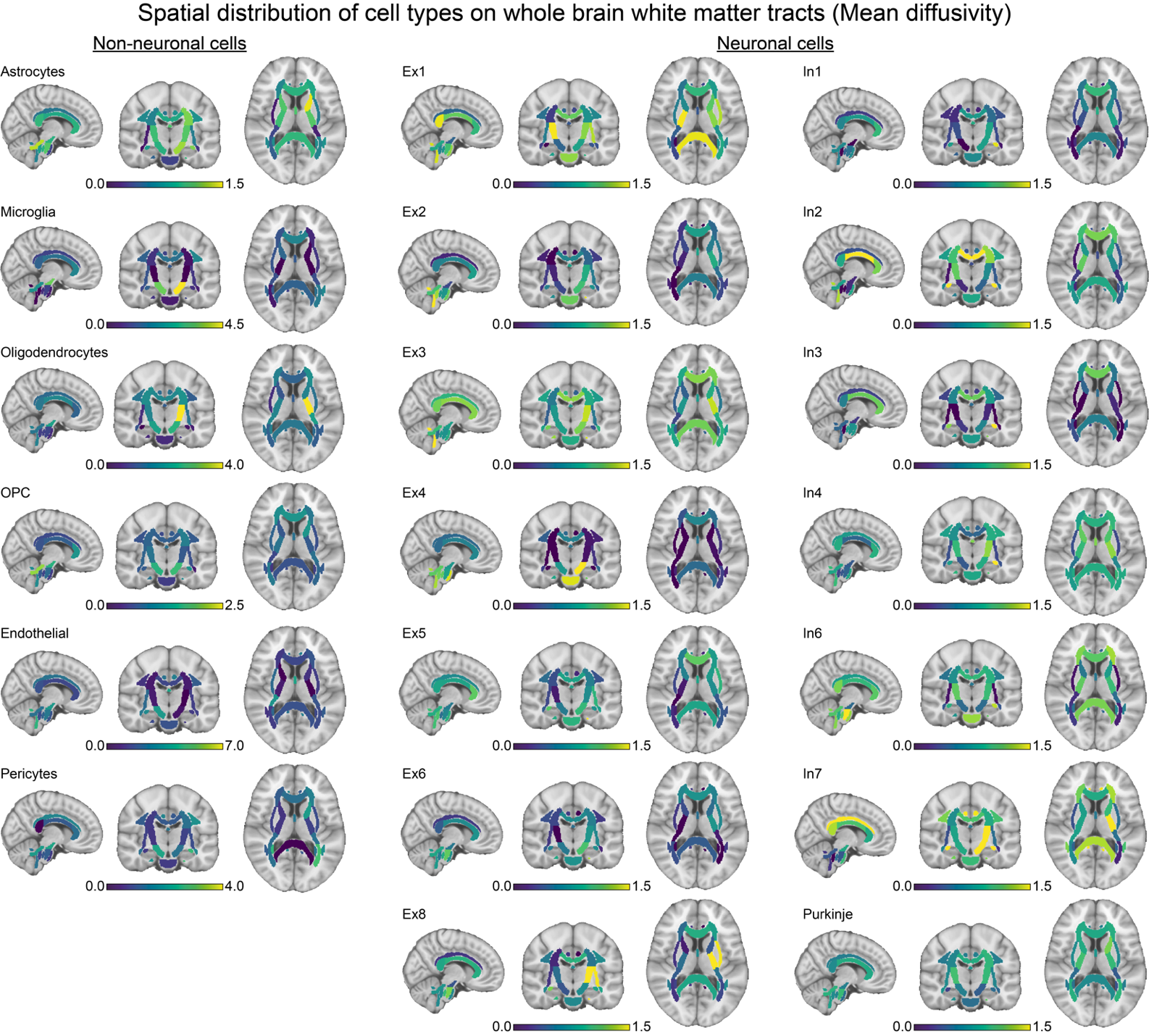

**Fig. S4.** Spatial distribution of cell type enrichment on whole brain white matter tracts as measured by mean diffusivity (MD). Individual cell type enrichment distributions across twenty cell subtypes are sectioned into neuronal (right panel) and non-neuronal groups (left panel) and displayed accordingly. Dark blue colors indicate low enrichment, while bright yellow indicates high enrichment.

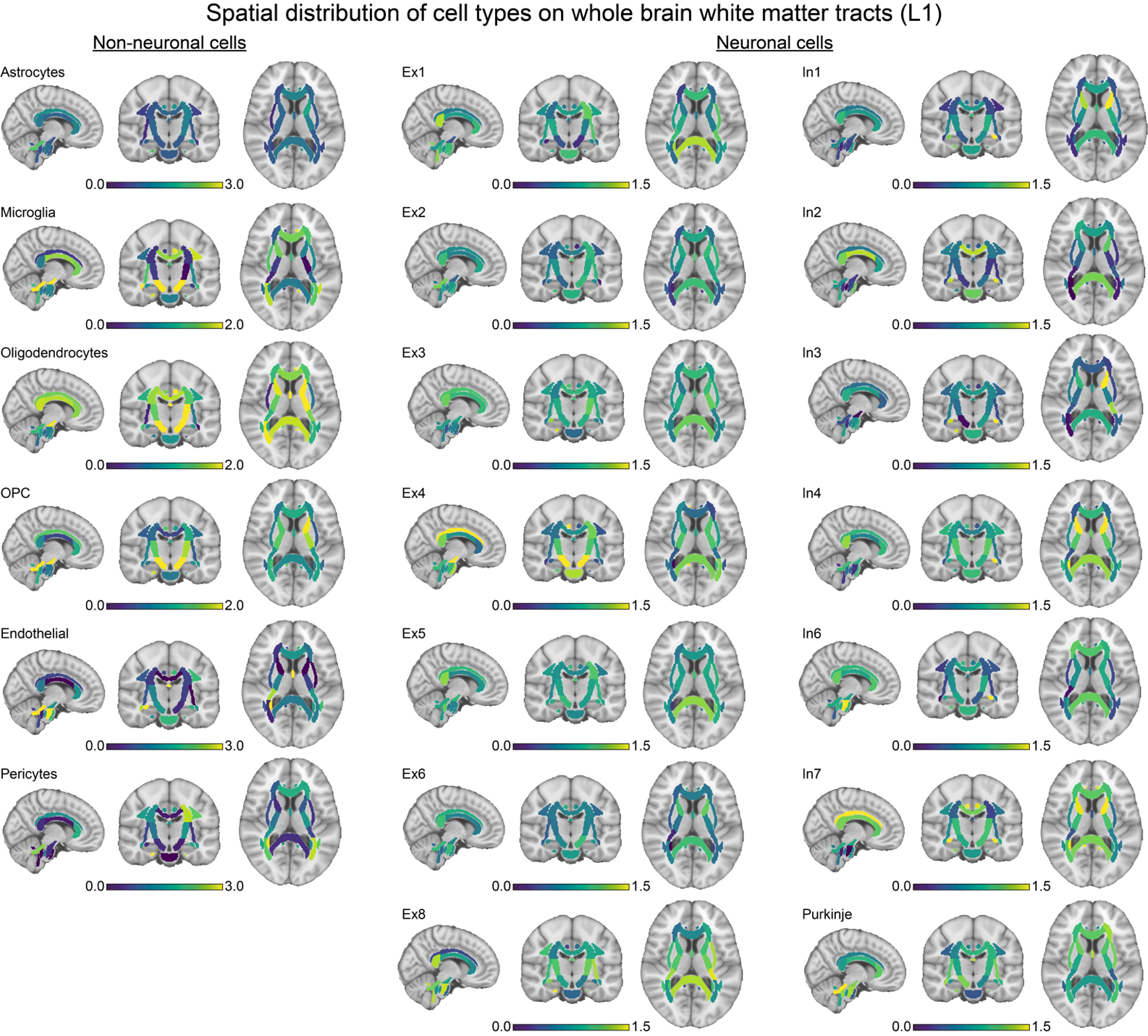

**Fig. S5.** Spatial distribution of cell type enrichment on whole brain white matter tracts as measured by L1. Individual cell type enrichment distributions across twenty cell subtypes are sectioned into neuronal (right panel) and non-neuronal groups (left panel) and displayed accordingly. Dark blue colors indicate low enrichment, while bright yellow indicates high enrichment.

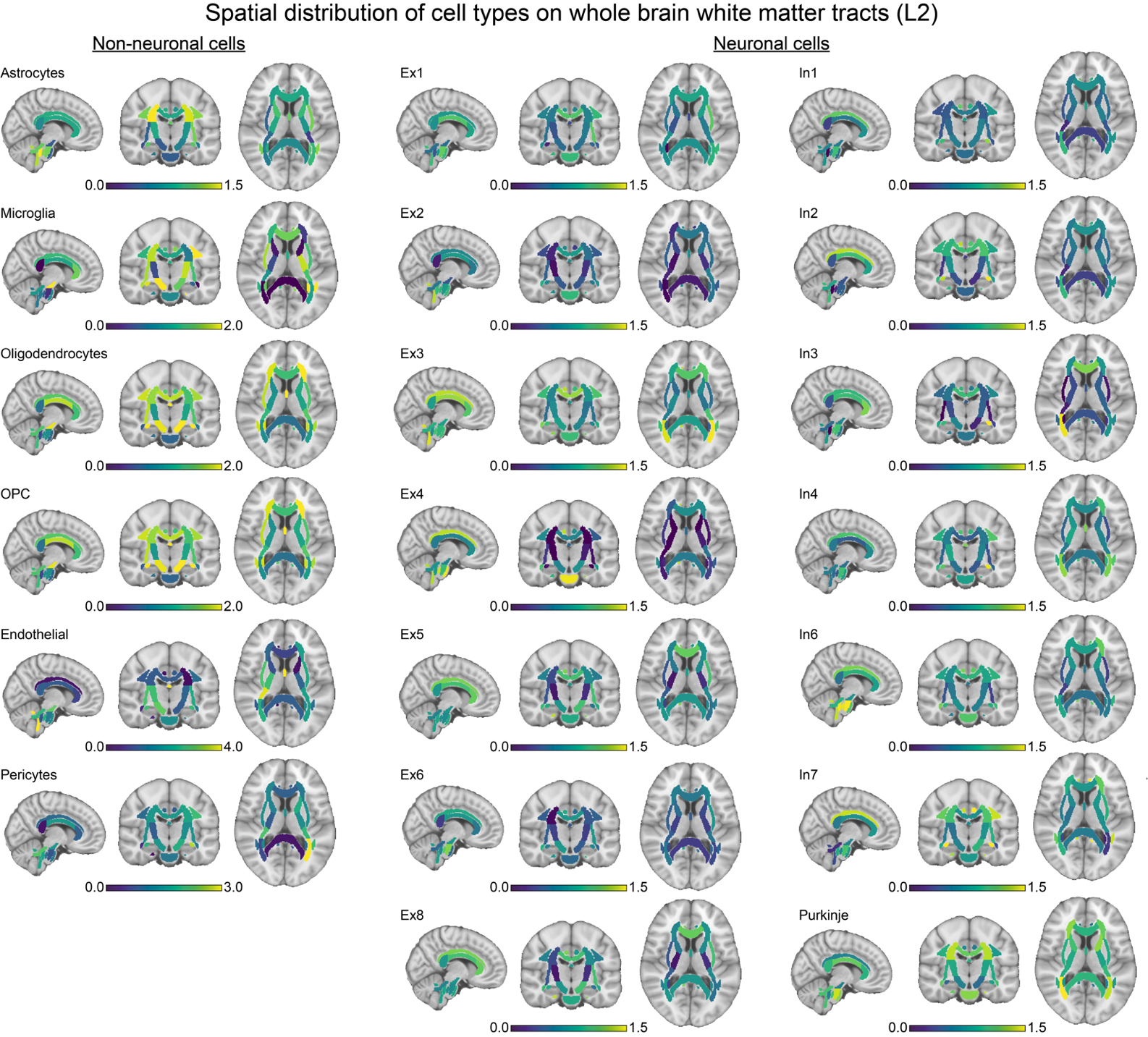

**Fig. S6.** Spatial distribution of cell type enrichment on whole brain white matter tracts as measured by L2. Individual cell type enrichment distributions across twenty cell subtypes are sectioned into neuronal (right panel) and non-neuronal groups (left panel) and displayed accordingly. Dark blue colors indicate low enrichment, while bright yellow indicates high enrichment.

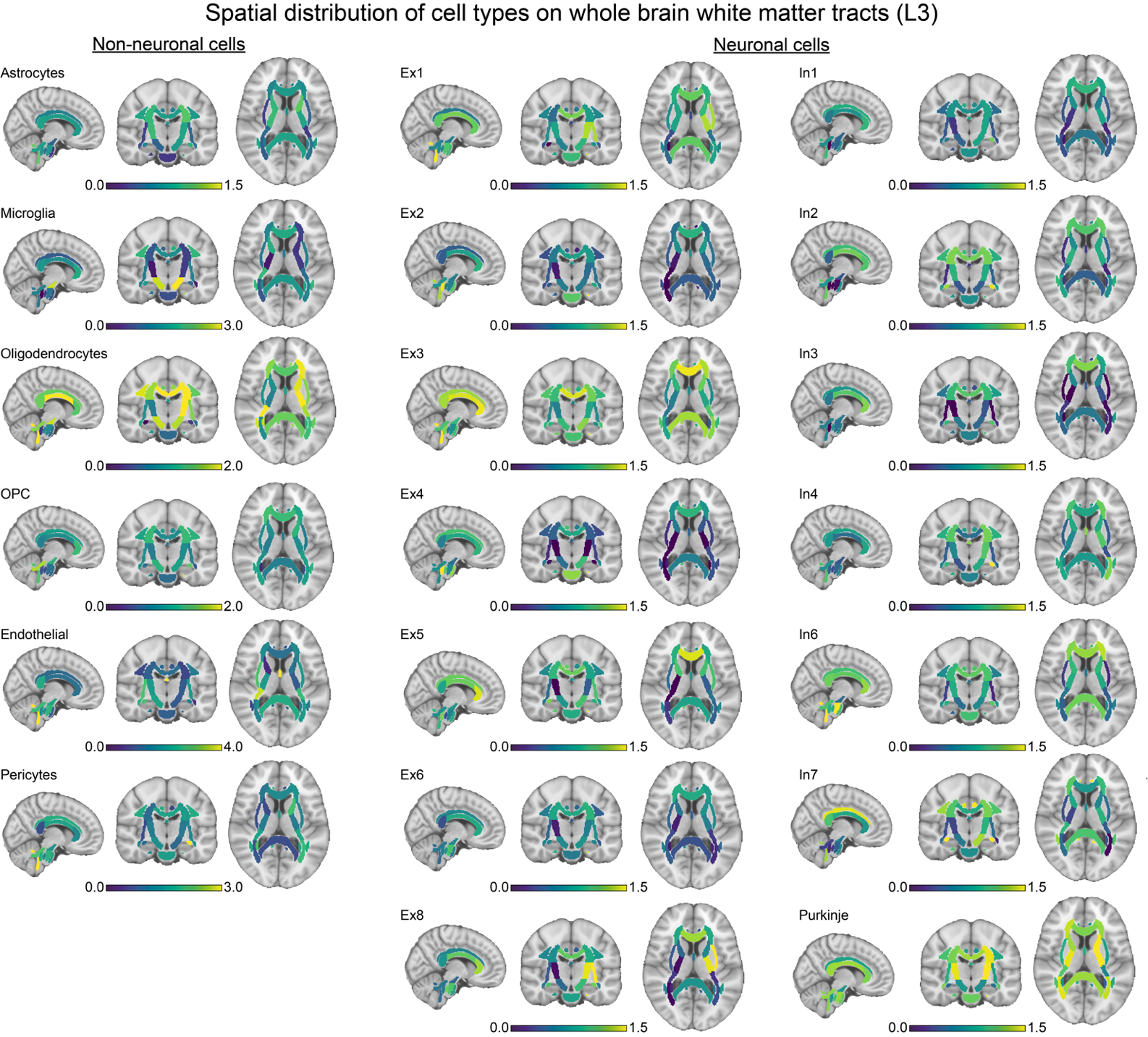

**Fig. S7.** Spatial distribution of cell type enrichment on whole brain white matter tracts as measured by L3. Individual cell type enrichment distributions across twenty cell subtypes are sectioned into neuronal (right panel) and non-neuronal groups (left panel) and displayed accordingly. Dark blue colors indicate low enrichment, while bright yellow indicates high enrichment.

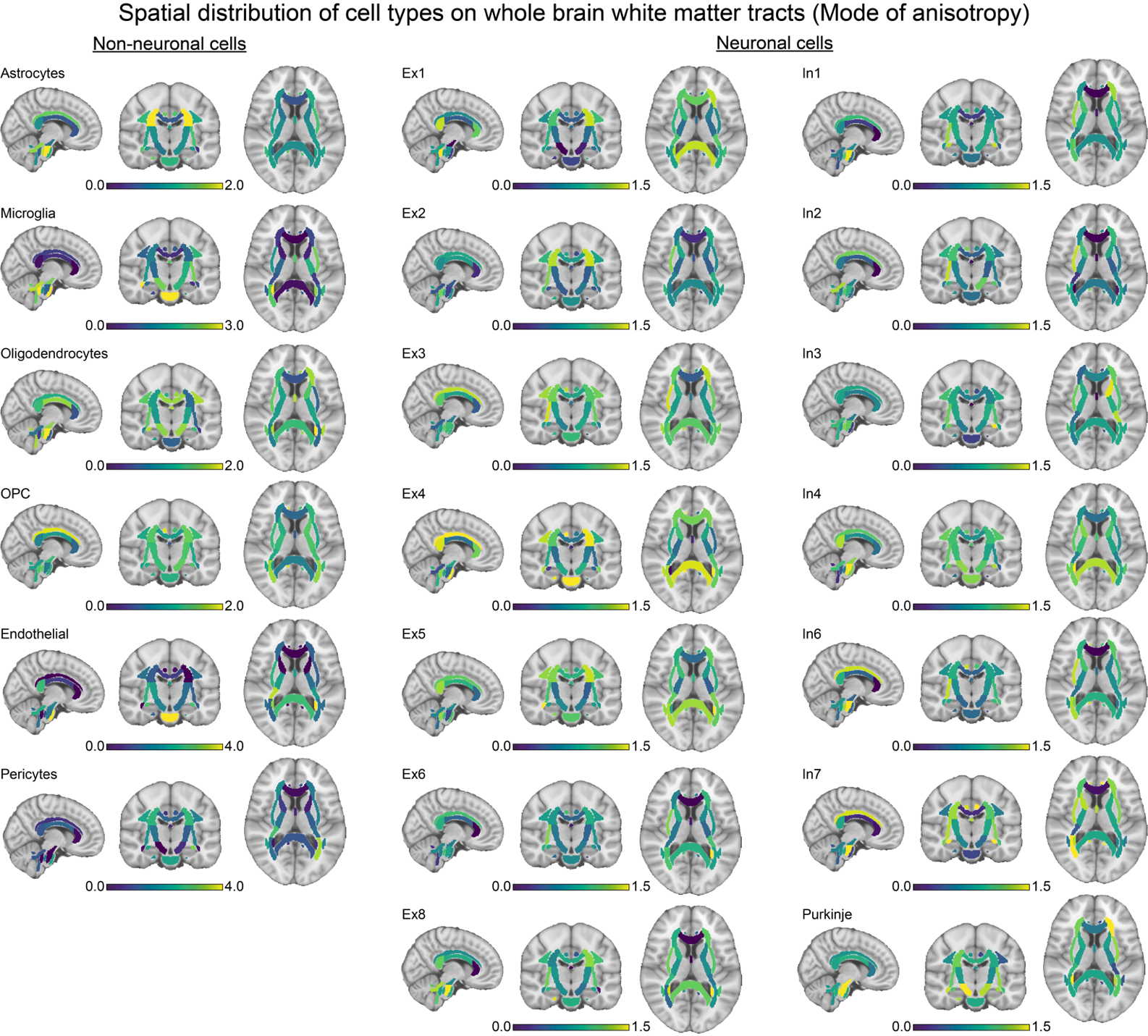

**Fig. S8.** Spatial distribution of cell type enrichment on whole brain white matter tracts as measured by mode of anisotropy (MO). Individual cell type enrichment distributions across twenty cell subtypes are sectioned into neuronal (right panel) and non-neuronal groups (left panel) and displayed accordingly. Dark blue colors indicate low enrichment, while bright yellow indicates high enrichment.

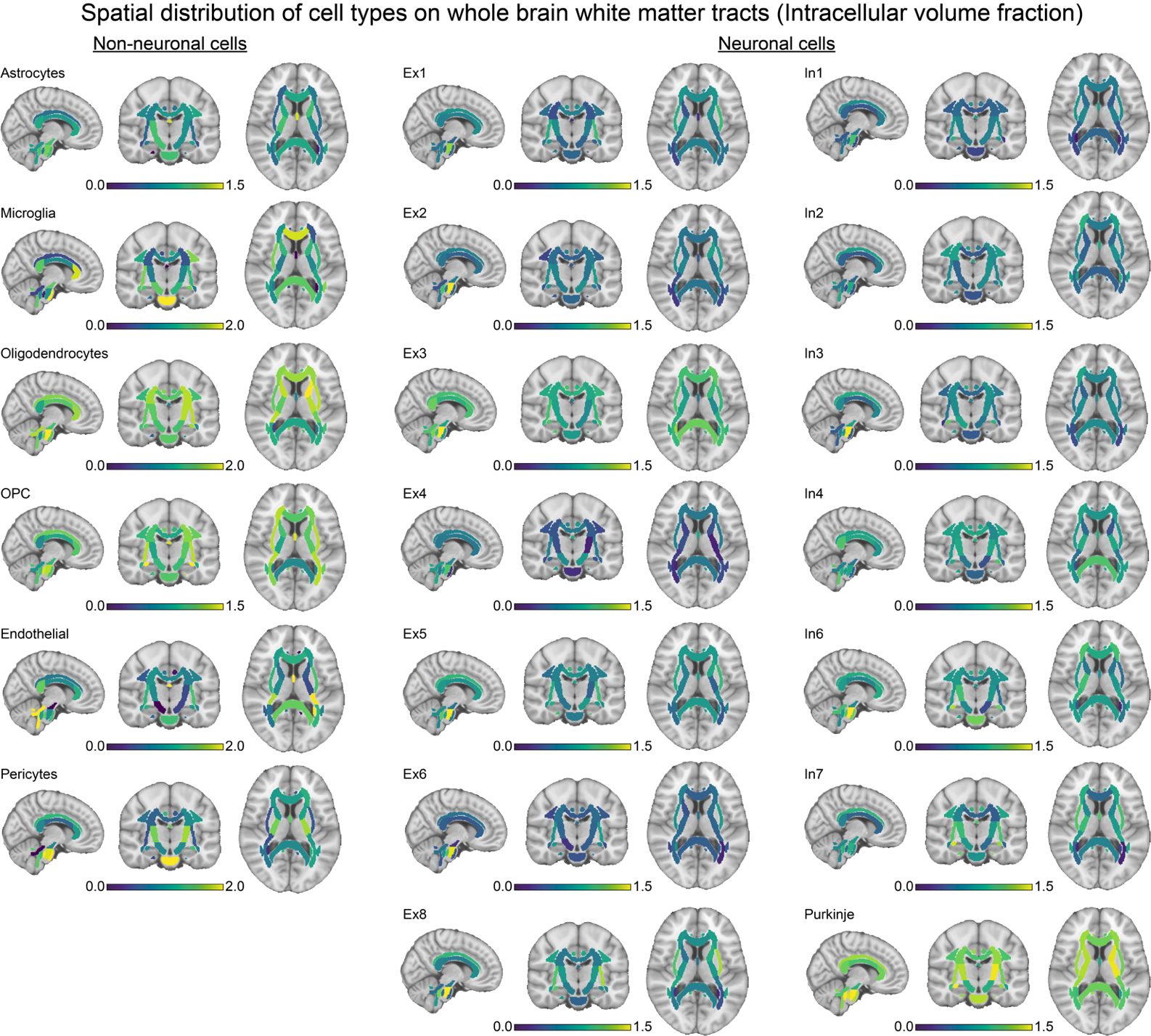

**Fig. S9.** Spatial distribution of cell type enrichment on whole brain white matter tracts as measured by intracellular volume fraction (ICVF). Individual cell type enrichment distributions across twenty cell subtypes are sectioned into neuronal (right panel) and non-neuronal groups (left panel) and displayed accordingly. Dark blue colors indicate low enrichment, while bright yellow indicates high enrichment.

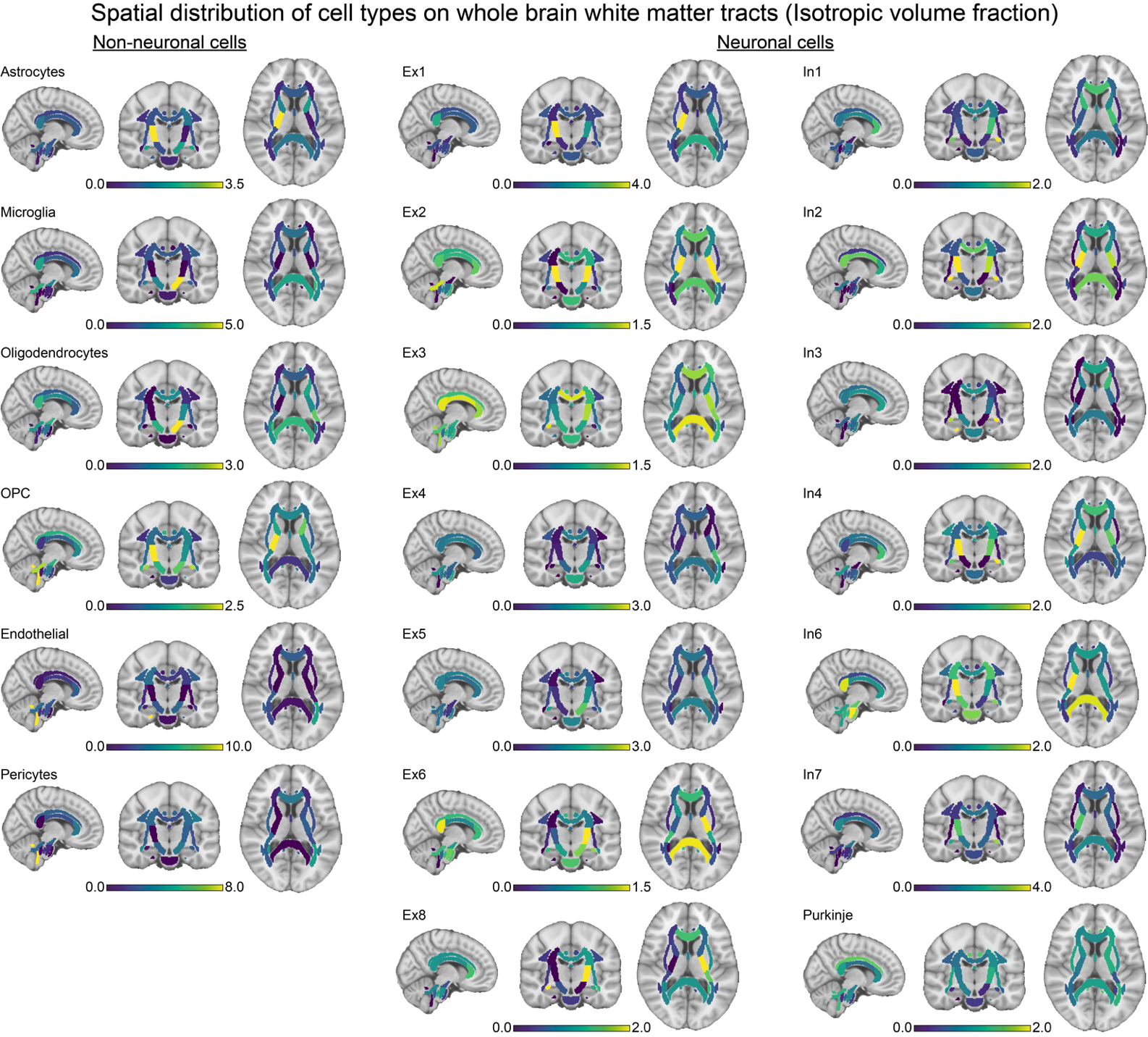

**Fig. S10.** Spatial distribution of cell type enrichment on whole brain white matter tracts as measured by isotropic volume fraction (ISOVF). Individual cell type enrichment distributions across twenty cell subtypes are sectioned into neuronal (right panel) and non-neuronal groups (left panel) and displayed accordingly. Dark blue colors indicate low enrichment, while bright yellow indicates high enrichment.

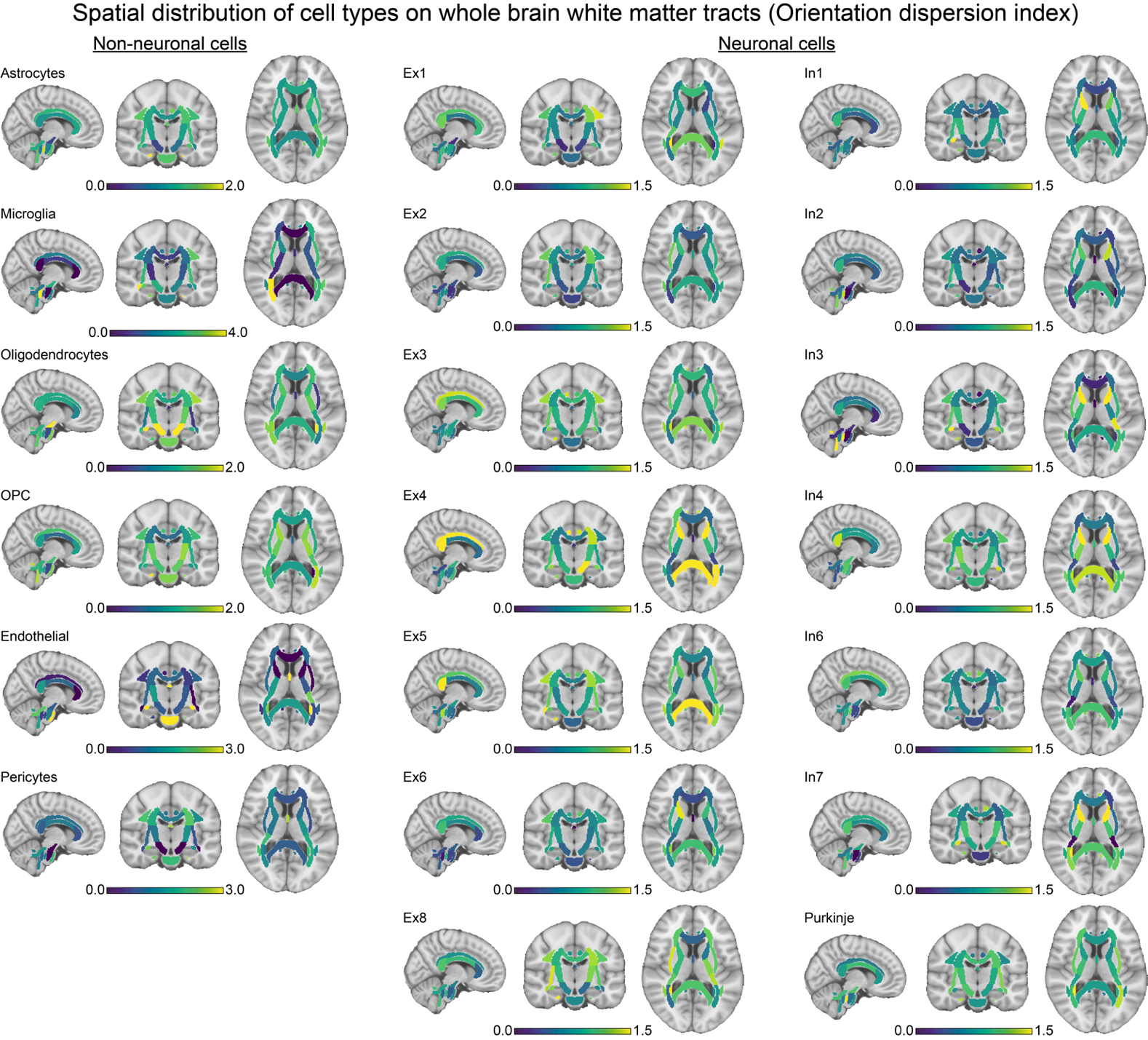

**Fig. S11.** Spatial distribution of cell type enrichment on whole brain white matter tracts as measured by orientation dispersion index (OD). Individual cell type enrichment distributions across twenty cell subtypes are sectioned into neuronal (right panel) and non-neuronal groups (left panel) and displayed accordingly. Dark blue colors indicate low enrichment, while bright yellow indicates high enrichment.

**Table S1.** Overview of white matter tracts derived from tract-based spatial statistics (TBSS) analyses mapped based on JHU ICBMB-DTI-81 white matter atlas and their respective abbreviations.

| **Abbreviation** | **White Matter Tract** |
| --- | --- |
| ACR | Anterior corona radiata |
| AIC | Anterior limb of internal capsule |
| BodyCC | Body of corpus callosum |
| CingCG | Cingulum, cingulate gyrus |
| CingHipp | Cingulum, hippocampus |
| CP | Cerebral peduncle |
| CST | Corticospinal tract |
| EC | External capsule |
| Fornix | Fornix |
| FornixST | Fornix cres and stria terminalis |
| GenuCC | Genu of corpus callosum |
| ICP | Inferior cerebellar peduncle |
| MCP | Middle cerebellar peduncle |
| ML | Medial lemniscus |
| PCR | Posterior corona radiata |
| PCT | Pontine crossing tract |
| PIC | Posterior limb of internal capsule |
| PTR | Posterior thalamic radiation |
| RIC | Retrolenticular part of internal capsule |
| SCP | Superior cerebellar peduncle |
| SCR | Superior coronal radiata |
| SFOF | Superior fronto-occipital fasciculus |
| SLF | Superior longitudinal fasciculus |
| SpleniumCC | Splenium of the corpus callosum |
| SS | Sagittal stratum |
| Tapetum | Tapetum |
| UF | Uncinate fasciculus |

**Table S2.** Classification of 48 white matter tracts derived from tract-based spatial statistics (TBSS) analyses mapped based on JHU ICBMB-DTI-81 white matter atlas.

| **White Matter Tract Class Classification** | |
| --- | --- |
| **Tract Class** | **White Matter Tracts** |
| Association | Cingulum – Hippocampus, Cingulum gyrus, Fornix, Fornix-stria terminalis, External capsule, Sagittal Striatum, Superior longitudinal fasciculus, Superior frontal-occipital fasciculus, Uncinate fasciculus |
| Commissural | Corpus callosum – body, genu, splenium, Pontine crossing tract, Tapetum |
| Projection | Corticospinal tract, Medial Lemniscus, Internal capsule – anterior, retrolenticular, posterior limbs, Corona radiata – anterior, posterior superior, Posterior thalamic radiation |
| Complex (Cerebellar) | Cerebral peduncle, Cerebellar peduncle – Superior, Middle, Inferior |

**Table S3.** Tract class comparisons of bivariate genetic correlation scores (r_g_) between 48 pairs of white matter tracts for each diffusivity measure, repeated across the 9 diffusivity measures. Within- and between- tract class differences for genetic correlation estimates (r_g_) were tested using paired sample t-tests. Table cells display resultant t-values and p-values respectively.

| **Tract Class Comparison** | **DMRI measure** | | | | | | | | |
| --- | --- | --- | --- | --- | --- | --- | --- | --- | --- |
|  | **FA** | **MD** | **L1** | **L2** | **L3** | **MO** | **ICVF** | **ISOVF** | **OD** |
| Association vs all others | t=2.36,  p=0.019 | t=5.89,  p=1.779e-8 | t=6.98, p=5.992e-11 | t=1.91, p=0.057 | t=1.91, p=0.057 | t=3.12, p=0.002 | t=6.99, p=4.975e-11 | t=4.19, p=4.203e-5 | t=5.44, p=1.888e-7 |
| Commissural vs all others | t=2.68, p=0.020 | t=0.53  p=0.605 | t=0.25, p=0.810 | t=-0.89 p=0.391 | t=-0.89, p=0.391 | t=-0.52, p=0.615 | t=1.12, p=0.285 | t=2.20, p=0.047 | t=2.24, p=0.044 |
| Complex vs all others | t=2.78,  p=0.011 | t=4.27,  p=3.265e-4 | t=3.72,  p=0.001 | t=1.90, p=0.071 | t=1.91, p=0.057 | t=3.59, p=0.002 | t=2.29, p=0.033 | t=5.65, p=1.347e-5 | t=4.20, p=4.165e-4 |
| Projection vs all others | t=-13.37, p<2.2e-16 | t=-3.62,  p=3.834e-4 | t=-4.23,  p=3.627e-5 | t=-7.60, p=9.35e-13 | t=-4.31, p= 2.532e-05 | t=-3.36,  p= 9.519e-4 | t=-6.42,  p=1.017e-9 | t=-8.79, p=1.433e-15 | t=-3.36, p=9.432e-4 |

**Table S4.** Univariate cellular associates of age-regressed white matter tract estimates. Table cells display correlations between age-regressed estimates and enrichment of twenty cell types, calculated by Spearman’s Rho, p_FDR_ reflects associated p-values after multiple comparison correction (FDR; Benjamini & Hochberg).

| **Cell**  **Type** | **FA** | | **MD** | | **L1** | | **L2** | | **L3** | | **MO** | | **ICVF** | | **ISOVF** | | **OD** | |
| --- | --- | --- | --- | --- | --- | --- | --- | --- | --- | --- | --- | --- | --- | --- | --- | --- | --- | --- |
|  | r | p_FDR_ | r | p_FDR_ | r | p_FDR_ | r | p_FDR_ | r | p_FDR_ | r | p_FDR_ | r | p_FDR_ | r | p_FDR_ | r | p_FDR_ |
| Ast | -0.065 | 0.697 | 0.049 | 0.870 | 0.103 | 0.647 | -0.179 | 0.399 | 0.191 | 0.426 | 0.068 | 0.810 | 0.244 | 0.327 | 0.386 | 0.058 | 0.123 | 0.808 |
| Mic | 0.009 | 0.951 | 0.117 | 0.608 | -0.007 | 0.964 | 0.204 | 0.365 | 0.106 | 0.632 | -0.023 | 0.877 | -0.087 | 0.619 | 0.426 | 0.031 | 0.316 | 0.175 |
| Oli | -0.552 | 0.001 | 0.133 | 0.568 | 0.399 | 0.027 | 0.522 | 0.003 | 0.123 | 0.620 | -0.150 | 0.789 | -0.175 | 0.367 | 0.076 | 0.823 | -0.059 | 0.918 |
| OPC | -0.099 | 0.556 | 0.031 | 0.870 | 0.178 | 0.374 | 0.173 | 0.399 | 0.052 | 0.811 | -0.106 | 0.789 | -0.400 | 0.327 | 0.191 | 0.382 | 0.072 | 0.918 |
| End | -0.183 | 0.481 | -0.183 | 0.474 | -0.028 | 0.942 | 0.107 | 0.583 | -0.164 | 0.529 | 0.040 | 0.810 | 0.274 | 0.327 | -0.047 | 0.984 | -0.047 | 0.918 |
| Per | 0.255 | 0.178 | -0.183 | 0.474 | 0.202 | 0.361 | 0.143 | 0.473 | -0.288 | 0.147 | -0.125 | 0.789 | 0.148 | 0.393 | 0.086 | 0.806 | -0.068 | 0.918 |
| Pur | 0.219 | 0.268 | 0.039 | 0.870 | 0.159 | 0.431 | -0.072 | 0.690 | -0.007 | 0.965 | -0.116 | 0.789 | -0.178 | 0.367 | -0.107 | 0.777 | -0.060 | 0.918 |
| Ex1 | -0.179 | 0.318 | -0.099 | 0.669 | 0.222 | 0.361 | 0.166 | 0.399 | 0.115 | 0.622 | 0.240 | 0.789 | -0.214 | 0.327 | 0.138 | 0.639 | -0.181 | 0.588 |
| Ex2 | -0.212 | 0.268 | -0.333 | 0.106 | 0.200 | 0.361 | -0.004 | 0.976 | -0.058 | 0.811 | 0.124 | 0.789 | -0.031 | 0.832 | -0.060 | 0.828 | -0.164 | 0.588 |
| Ex3 | -0.157 | 0.382 | -0.132 | 0.568 | 0.467 | 0.009 | 0.118 | 0.563 | -0.024 | 0.918 | -0.047 | 0.810 | 0.205 | 0.327 | -0.055 | 0.841 | -0.171 | 0.588 |
| Ex4 | -0.124 | 0.481 | -0.519 | 0.004 | -0.012 | 0.964 | -0.078 | 0.690 | -0.283 | 0.147 | 0.188 | 0.789 | 0.103 | 0.568 | -0.203 | 0.382 | 0.020 | 0.924 |
| Ex5 | -0.339 | 0.051 | -0.024 | 0.870 | 0.182 | 0.374 | 0.336 | 0.085 | 0.337 | 0.134 | 0.114 | 0.789 | -0.210 | 0.327 | -0.196 | 0.382 | -0.237 | 0.527 |
| Ex6 | -0.373 | 0.032 | -0.356 | 0.106 | 0.050 | 0.867 | 0.170 | 0.399 | 0.139 | 0.620 | 0.089 | 0.810 | -0.154 | 0.393 | -0.255 | 0.232 | -0.366 | 0.112 |
| Ex8 | -0.335 | 0.051 | -0.149 | 0.568 | 0.259 | 0.302 | 0.284 | 0.146 | 0.131 | 0.620 | 0.206 | 0.789 | -0.224 | 0.327 | 0.002 | 0.999 | -0.365 | 0.112 |
| In1 | -0.552 | 0.001 | 0.272 | 0.183 | 0.417 | 0.023 | 0.391 | 0.058 | 0.336 | 0.134 | 0.058 | 0.810 | -0.311 | 0.327 | 0.284 | 0.172 | 0.181 | 0.588 |
| In2 | -0.397 | 0.027 | 0.270 | 0.183 | 0.196 | 0.361 | 0.376 | 0.058 | 0.295 | 0.147 | -0.119 | 0.789 | -0.167 | 0.367 | 0.380 | 0.058 | 0.031 | 0.924 |
| In3 | -0.387 | 0.028 | 0.036 | 0.870 | 0.072 | 0.784 | 0.232 | 0.281 | 0.232 | 0.280 | 0.061 | 0.810 | -0.204 | 0.327 | 0.113 | 0.813 | 0.015 | 0.924 |
| In4 | -0.462 | 0.007 | 0.319 | 0.110 | 0.574 | 0.001 | 0.333 | 0.085 | 0.439 | 0.039 | 0.118 | 0.789 | -0.060 | 0.719 | 0.284 | 0.172 | -0.041 | 0.918 |
| In6 | -0.183 | 0.318 | -0.141 | 0.568 | -0.151 | 0.434 | -0.066 | 0.690 | 0.051 | 0.811 | -0.066 | 0.810 | 0.172 | 0.367 | 0.010 | 0.995 | 0.102 | 0.888 |
| In7 | -0.183 | 0.318 | 0.342 | 0.106 | 0.246 | 0.305 | 0.300 | 0.128 | 0.284 | 0.147 | -0.189 | 0.789 | -0.256 | 0.327 | 0.226 | 0.172 | 0.220 | 0.529 |
